# Uncovering the isoform-resolution kinetic landscape of nonsense-mediated mRNA decay with EZbakR

**DOI:** 10.1101/2025.03.12.642874

**Authors:** Justin W. Mabin, Isaac W. Vock, Martin Machyna, Nazmul Haque, Poonam Thakran, Alexandra Zhang, Ganesha Rai, Isabelle Nathalie-Marie Leibler, James Inglese, Matthew D. Simon, J. Robert Hogg

**Author notes:** Paul-Ehrlich-Institut, Host-Pathogen-Interactions, 63225 Langen, Germany (Martin Machyna). Ultragenyx, 7000 Shoreline Ct, South San Francisco, CA 94080. Equal contribution (Justin W. Mabin, Isaac W. Vock).

## Abstract

Cellular RNA levels are a product of synthesis and degradation kinetics, which can differ among transcripts of the same gene. An important cause of isoform-specific decay is the nonsense-mediated mRNA decay (NMD) pathway, which degrades transcripts with premature termination codons (PTCs) and other features. Understanding NMD functions requires strategies to quantify isoform kinetics; however, current approaches remain limited. Methods like nucleotide-recoding RNA-seq (NR-seq) enable insights into RNA kinetics, but existing bioinformatic tools do not provide robust, isoform-specific degradation rate constant estimates. We extend the EZbakR-suite by implementing a strategy to infer isoform-level kinetics from short-read NR-seq data. This approach uncovers unexpected variability in NMD efficiency among transcripts with conserved PTC-containing exons and rapid decay of a subset of mRNAs lacking PTCs. Our findings highlight the effects of competition between NMD and other decay pathways, provide mechanistic insights into established NMD efficiency correlates, and identify transcript features promoting efficient decay.

## Background

Transcript isoform abundance reflects the balance between the rates of RNA synthesis and degradation. Alternative splicing is a well-known mechanism for modulating isoform expression, however the degree to which differential turnover kinetics contribute to isoform-level regulation remains less explored.

A prominent example of degradation-driven transcript isoform regulation is nonsense-mediated mRNA decay (NMD). NMD is a conserved translation-dependent RNA turnover mechanism with essential quality control and regulatory functions. The NMD pathway recognizes and degrades mRNA isoforms containing premature translation termination codons (PTCs), which can arise from genetic mutations or errors in mRNA biogenesis [1,2]. Consequently, NMD is a central modulator of the phenotypic consequences of nonsense mutations in cancers and genetic disease [3]. Beyond quality control, NMD plays a major role in cellular gene expression regulation. Endogenous NMD substrates are frequently PTC-containing transcripts generated by alternative splicing but also include mRNAs with upstream open reading frames (uORFs) or long 3’ untranslated regions (3’UTRs) [4].

Much is still uncertain about the features that determine whether and how efficiently a transcript is degraded by NMD. The presence of an intron more than 50 nucleotides downstream of the termination codon is a strong predictor of NMD susceptibility [5]. Transcripts violating this rule are detected by NMD due to the persistence of exon junction complexes (EJCs) on their 3’UTRs, which enhances NMD factor association [6,7]. Despite the utility of the 50-nucleotide rule, current models struggle to explain variable decay of transcripts predicted to contain PTCs. Studies of PTCs in paired genomic and transcriptomic datasets from tumors and healthy tissues have identified correlates of low NMD efficiency, such as a short distance between the mRNA 5’ end and the PTC or a long distance between the PTC and the next downstream exon junction [8–10]. Still, a substantial fraction (≥30%) of variability in decay of putative PTC-containing mRNAs is unexplained [11], with even greater uncertainty surrounding how decay specificity and efficiency are determined for NMD targets lacking PTCs [12].

The lack of effective methods for quantifying the turnover kinetics of individual transcript isoforms has been an important technical barrier to investigations of NMD. Consequently, most studies of NMD have instead relied on differential transcript abundance as a proxy for NMD activity, which can be confounded by indirect transcriptional upregulation in response to NMD perturbation. In addition, transcriptome-wide measurements of transcript degradation are required to understand how competition between NMD and other decay pathways affect quality control and gene expression regulation. Therefore, new strategies are needed to directly probe isoform degradation kinetics.

Nucleotide-recoding RNA-seq (NR-seq) methods such as TimeLapse-seq or SLAM-seq, provide a promising experimental route by which to overcome this challenge [13,14]. NR-seq uses metabolic labeling and nucleotide recoding chemistry to probe the synthesis and degradation kinetics of mRNAs without having to globally inhibit transcription [13–15]. NR-seq also does not require biochemical enrichment of labeled RNA, circumventing challenges of unlabeled RNA contamination and normalization across input and enrichment samples [16]. However, existing bioinformatic tools are not equipped to perform kinetic parameter estimation with isoform resolution. We have recently developed the EZbakR-suite, which improves and generalizes many aspects of NR-seq data processing and analysis [17]. In particular, the EZbakR-suite includes a pipeline (fastq2EZbakR) for flexibly assigning sequencing reads to different feature sets, such as transcript equivalence classes (TECs, the set of isoforms with which a read is compatible). This read assignment strategy facilitates estimating transcript isoform kinetic parameters from NR-seq data.

Here, we extend the functionality of the EZbakR-suite by implementing a novel method to infer isoform-specific kinetic parameters. Analyses of simulated and real data confirm that this approach provides accurate estimates and captures well-established features of RNA turnover dynamics. We further implement a strategy for refinement and filtering of annotations, which we find to be important for accurate quantification of transcript degradation rate constants. To showcase the power of this approach, we use the EZbakR-suite in combination with acute NMD inhibition to map the kinetic landscape of human NMD. We quantify transcript degradation rate constants in the presence and absence of a functional NMD pathway, revealing the influence of NMD on global mRNA decay kinetics. Our data identify previously unappreciated variability in the decay kinetics of transcripts containing highly conserved poison exons and highlight the influence of competing RNA decay pathways on both the regulatory and quality control functions of NMD. We additionally identify a set of target mRNAs that lack PTCs but nevertheless are subject to highly efficient NMD. Together, our data provide mechanistic explanations for previously identified correlates of NMD efficiency, identify transcript features associated with efficient decay, and provide a roadmap for future studies of transcript isoform turnover regulation. This work illustrates how isoform-level analyses and the EZbakR-suite can be used to deepen our understanding of RNA kinetic regulation.

## Results

### Transcript isoform kinetic parameter estimation

RNA biogenesis and decay pathways often differentially affect transcript isoforms produced from the same gene. Accurate studies of RNA synthesis and decay mechanisms therefore require the ability to measure isoform-level kinetics. To meet this need, we developed a strategy to infer transcript isoform synthesis and degradation rate constants from short read NR-seq data (**Figure 1A**). This strategy is implemented as a part of our recently released EZbakR-suite, which includes a Snakemake preprocessing pipeline (fastq2EZbakR) and an NR-seq analysis R package (EZbakR) [17].

**Figure 1.**
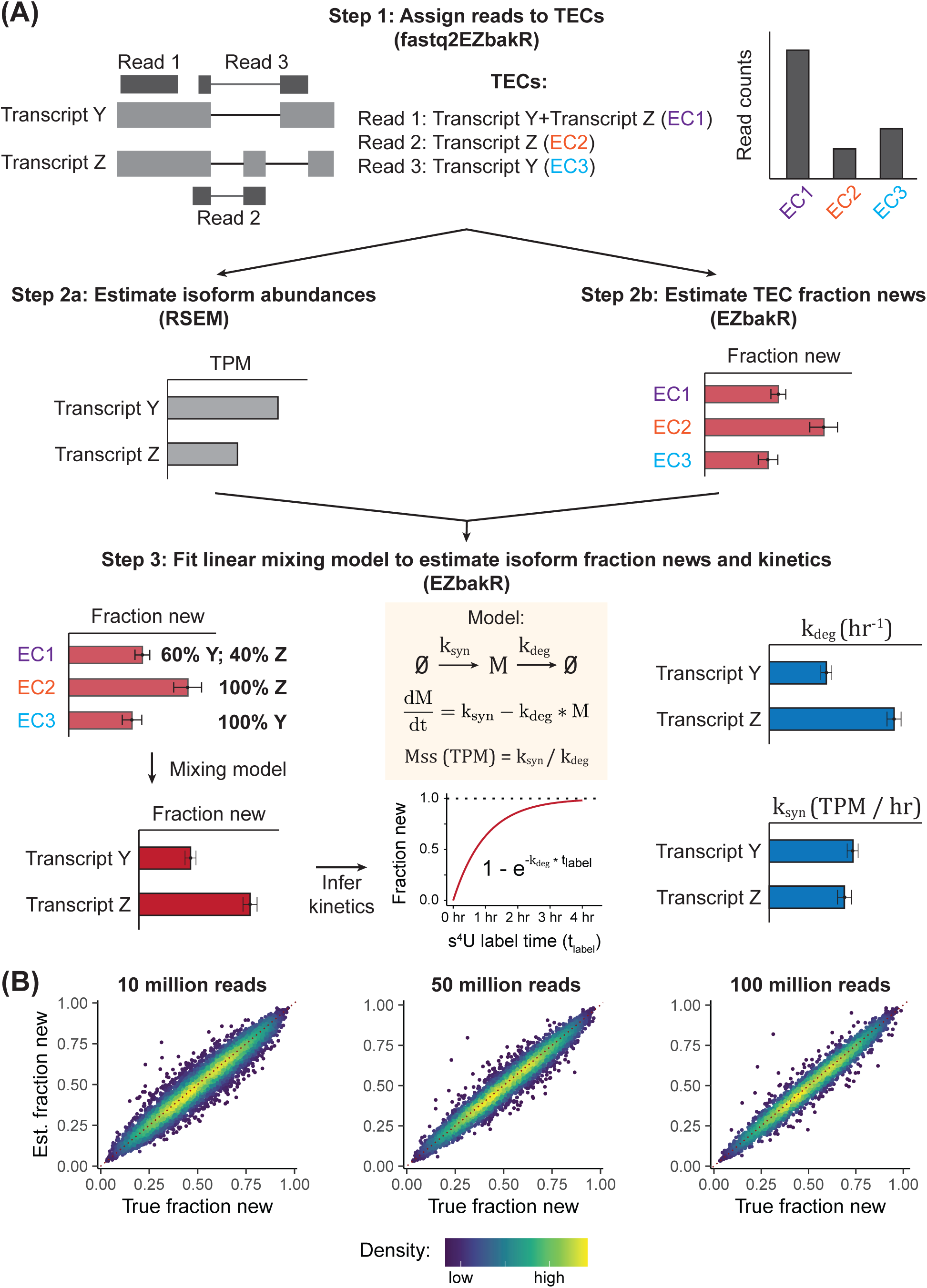
Isoform-level NR-seq analyses. **(A)** Schematic of the EZbakR-suite’s isoform analysis pipeline. **Top:** fastq2EZbakR parses transcriptome alignment files generated by STAR to assign sequencing reads to their transcript equivalence class (TEC). **Middle:** Transcript isoform quantification is performed by RSEM within fastq2EZbakR, and the TEC fraction of reads from RNAs synthesized in the presence of metabolic label (TEC fraction new) is estimated by EZbakR. **Bottom**: Isoform abundance estimates are used to infer isoform-specific fraction news from the TEC fraction news via a linear mixing model. Isoform fraction news and abundances are converted to synthesis and degradation rate constants via a standard kinetic model of NR-seq data. **(B)** Simulated data assessment of isoform fraction new estimate accuracy across a range of read depths. Points are colored by density, and the red dotted line represents perfect estimation (y = x).

NR-seq allows identification of reads derived from transcripts produced during a metabolic labeling period, and the ratio of labeled (new) reads to total reads (the fraction new) can then be used to infer kinetic parameters [13,18]. Our strategy for isoform-level NR-seq analysis works by combining transcript isoform quantification using existing tools like RSEM with estimates of the fraction of reads from a given transcript equivalence class (TEC) that come from labeled RNA (i.e., the TEC fraction new) [19]. fastq2EZbakR implements TEC read assignment (**Figure 1A**, top), and EZbakR estimates fraction news for each TEC (**Figure 1A**, middle). We have added a novel linear mixing model to EZbakR, which combines the isoform abundance and TEC fraction new estimates to accurately infer transcript isoform fraction news (**Figure 1A**, bottom). From these, transcript isoform degradation rate constants are inferred as described previously (**Figure 1A**, bottom; see also Methods). Transcript isoform degradation rate constant estimates and abundance estimates can then be combined to infer synthesis rate constants. Analyses of simulated data confirmed that EZbakR is able to accurately infer isoform fraction news (and thus kinetic parameters) across a range of sequencing depths, read lengths, and metabolic labeling efficiencies (**Figure 1B** and **S1**). EZbakR’s isoform-level analysis also provides consistent kinetic parameter estimates across replicates, datasets, and human cell lines (**Figure S2A**). EZbakR is thus the first tool able to extract isoform-level kinetic insights from short-read total RNA NR-seq data.

### Isoform-level analyses capture signal missed by gene-level analyses

The NMD pathway targets transcripts with specific features for RNA decay. Genes often produce a mixture of NMD-sensitive and NMD-insensitive transcripts, generated through alternative splicing, polyadenylation, and other processes, suggesting that understanding NMD mechanisms and roles would benefit from isoform-level quantification of RNA synthesis and degradation [20]. We therefore applied EZbakR’s isoform-level analyses to the task of identifying targets of NMD. To do so, we combined NR-seq with acute NMD inhibition and used EZbakR to identify significantly stabilized transcript isoforms (**Figure 2A**). We chose to inhibit NMD with a specific inhibitor of the SMG1 kinase (SMG1i), which prevents the phosphorylation of the key NMD factor UPF1 and inhibits NMD [21]. Because this approach allows for analysis of RNA decay following a short period of NMD inhibition, we reasoned that it would primarily capture direct effects of blocking the pathway, a prediction supported by our initial gene-level analyses, which revealed 408 genes with significant RNA stabilization but only 6 with significant destabilization (**Figure 2B**, left).

**Figure 2.**
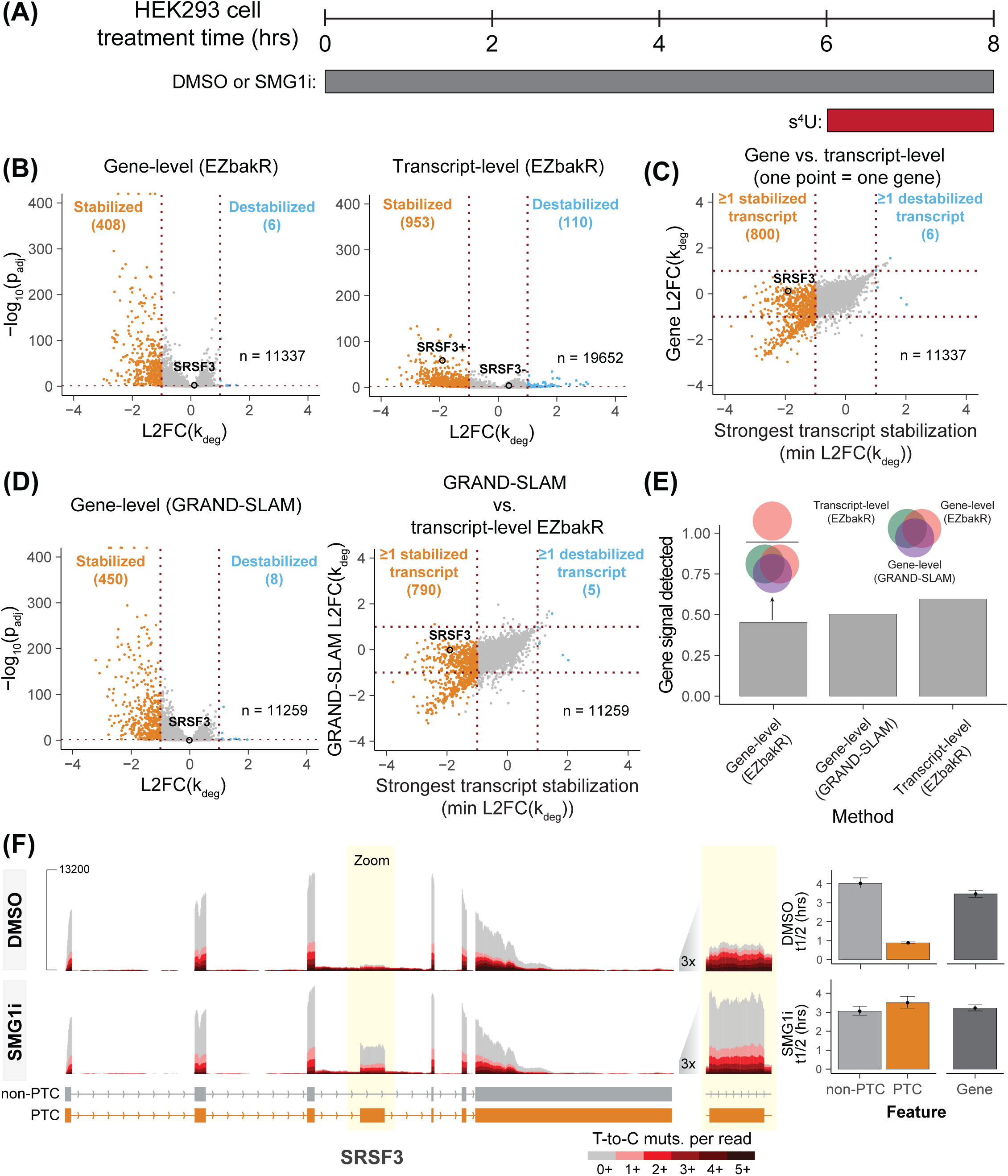
Isoform-level NR-seq analyses accurately identify targets of NMD. **(A)** Schematic for TimeLapse-seq experiment designed to identify targets of NMD. HEK293 cells were treated with either a SMG1 inhibitor (SMG1i) or DMSO for 8 hours. Cells were fed with s^4^U for the last 2 hours of the treatment. **(B)** Gene- and isoform-level analysis volcano plots. L2FC(k_deg_) is log_2_(SMG1i k_deg_ / DMSO k_deg_), i.e., the log_2_-fold change in the degradation rate constant estimate. Cutoff for significance is FDR < 0.01 and |L2FC(k_deg_)| > 1. For the gene-level analysis, only genes with at least 10 reads in all samples were included. For the isoform-level analyses, only isoforms with a TPM > 1 in all samples, as well as estimated read counts of 10 in all samples, were included. SRSF3+ refers to the PTC-containing isoform from the SRSF3 gene, while SRSF3-is the non-PTC isoform. **(C)** Direct comparison of gene- and isoform-level analyses. For each gene, the minimum isoform L2FC(k_deg_) (i.e., strongest stabilization) was compared to the gene-level L2FC(k_deg_). **(D)** Same as **B** left and **C**, but with gene-level analyses using GRAND-SLAM (multiple-test adjusted p-values calculated using EZbakR). **(E)** Comparison of the fraction of stabilization events identified by each analysis. A list of genes identified by any of the gene-level analyses or that produced at least one stabilized isoform was compiled. The fraction of these genes identified in each of the three analysis strategies tested was then calculated (y-axis). Venn diagrams schematize this scoring metric. **(F)** Example sequencing tracks of an NMD target identified by the isoform-level analysis but missed by the gene-level analysis (SRSF3). Tracks are colored by mutational content. Bar plots to the right of the tracks depict the estimated isoform-level and gene-level half-lives (t1/2), in hours.

To evaluate the benefits of EZbakR’s isoform-level analysis, we compared it to a standard gene-level analysis approach. We hypothesized that the isoform-level analysis would reveal NMD targets missed by gene-level analysis due to minor-isoform stabilization being obscured by data from unaffected isoforms. Both the isoform-level and gene-level analyses identified many stabilization events with high confidence (**Figure 2B**). Despite this, we noticed that several genes with NMD-sensitive transcripts identified in the isoform-level analysis were missed by the gene-level analysis. This included the well-established NMD-target producing gene SRSF3 [22,23], which we will use as a reference point throughout. Comparing the strongest isoform-level stabilization at each gene to the gene-level stabilization, we confirmed our hypothesis that while there was generally good agreement, a large number of gene-level effects were significantly smaller than the strongest isoform-level effect (middle-left section of **Figure 2C**). The same trend is observed when comparing the isoform-level EZbakR analysis to a gene-level analysis using a different NR-seq analysis tool, GRAND-SLAM (**Figure 2D**) [18]. To further explore this observation, we defined a “stabilized gene set” comprising genes identified as producing significantly stabilized RNA by the EZbakR-suite or GRAND-SLAM gene-level analyses or as producing at least one isoform identified as stabilized by the EZbakR-suite transcript-level analysis. Then from this comprehensive set of stabilization hits, we determined what fraction of these events were detected by each individual analysis (**Figure 2E**). This metric revealed that the isoform-level analysis was able to successfully identify more stabilization events than any of the gene-level analyses. The enhanced performance of the isoform-level analysis likely reflects NMD-sensitive isoforms comprising a small fraction of their gene’s total isoform pool, even after NMD inhibition (**Figure 2F**). Thus, EZbakR’s isoform-level analyses are able to capture NMD targets missed by a gene-level NR-seq analysis.

### NR-seq analysis uncovers signal invisible to RNA-seq

To assess the value of having isoform kinetic parameter estimates in addition to abundance estimates, we investigated the relationship between differential expression and differential stabilization in response to SMG1 inhibition. We found there to be a strong correlation between changes in isoform abundance and isoform degradation rate constants (**Figure S2B**, left). In particular, SMG1 inhibition resulted in a significant increase in both the abundance and stability of 554 isoforms. Despite this, there were 92 isoforms for which abundance significantly increased but there was little to no increase in stability (**Figure S2B**, right). These could represent indirect transcriptional effects of SMG1 inhibition that would have confounded a standard RNA-seq analysis. In addition, 151 isoforms appeared to be significantly stabilized while showing no increase in abundance. Some of these isoforms may represent instances of transcriptional repression in response to NMD inhibition that violates the steady-state assumption made by EZbakR, a challenge that can be curtailed with a slight modification to the NR-seq experimental design used here [24]. Alternatively, these may be instances of well-established homeostatic mechanisms that couple transcription and degradation so as to maintain RNA levels [25–27]. This analysis shows how NR-seq can improve identification of NMD targets.

NR-seq and EZbakR also allowed us to assess the extent to which the turnover kinetics of an isoform contribute to its abundance. We found isoform degradation rate constants to be strongly negatively correlated with both isoform abundance (**Figure S2C**, left) and the isoform fraction (the fraction of transcripts from a gene contributed by a specific isoform; **Figure S2C**, middle). To investigate this further, we quantified the coefficient of determination (R^2^; proportion of variance in data explained by a linear fit) for each of these relationships (**Figure S2C**, right). Isoform degradation rate constant differences explain around 38% of the isoform abundance variance, meaning that isoform turnover kinetics are a significant contributor to isoform expression levels. In addition, isoform degradation rate constant differences account for around 18% of the isoform fraction variance. This suggests not only that differences in the stability of isoforms from different genes are major determinants of the relative abundances of those isoforms, but also that differences in stabilities of isoforms from the same gene (and not just alternative splicing preferences) play a considerable role in establishing alternative isoform abundance. These analyses reveal that isoform turnover regulation is an important determinant of isoform expression levels.

### AnnotationCleaner: a pipeline for building custom annotations

While the EZbakR isoform analysis successfully identified many high-confidence NMD targets, we suspected that this analysis missed a number of unannotated NMD targets, as previous work has highlighted that numerous NMD targets are often absent from standard references (analyses in Figure 2 used the RefSeq hg38 annotation) [28–31]. In addition, recent work and our own observations indicated that discrepancies between references and the HEK293 transcriptome were leading to inaccuracies in the isoform-level analyses [32,33] **(Figure S3** and **S4A**). We therefore sought ways to build and stringently filter custom annotations.

Towards that end, we developed AnnotationCleaner, a pipeline to refine references with short and long read RNA-seq data. AnnotationCleaner combines reference-guided transcriptome assembly using the extensively validated tool StringTie with custom scripting to augment StringTie’s trimming of UTRs and flag poorly supported isoforms that should be filtered out in downstream analyses [34,35]; **Figure 3A**). Filtering was performed in two ways. First, we developed a new metric that identifies isoforms with low coverage (i.e., similar to pre-mRNA levels) of one or more exons and thus flags isoforms that do not have ample read coverage support in our RNA-seq data. Second, we used the previously described junction coverage compatibility (JCC) score (**Figure 3A**; see Methods for details) to flag entire genes whose read coverage distributions deviate from those predicted by the estimated relative isoform abundances [36,37].

**Figure 3.**
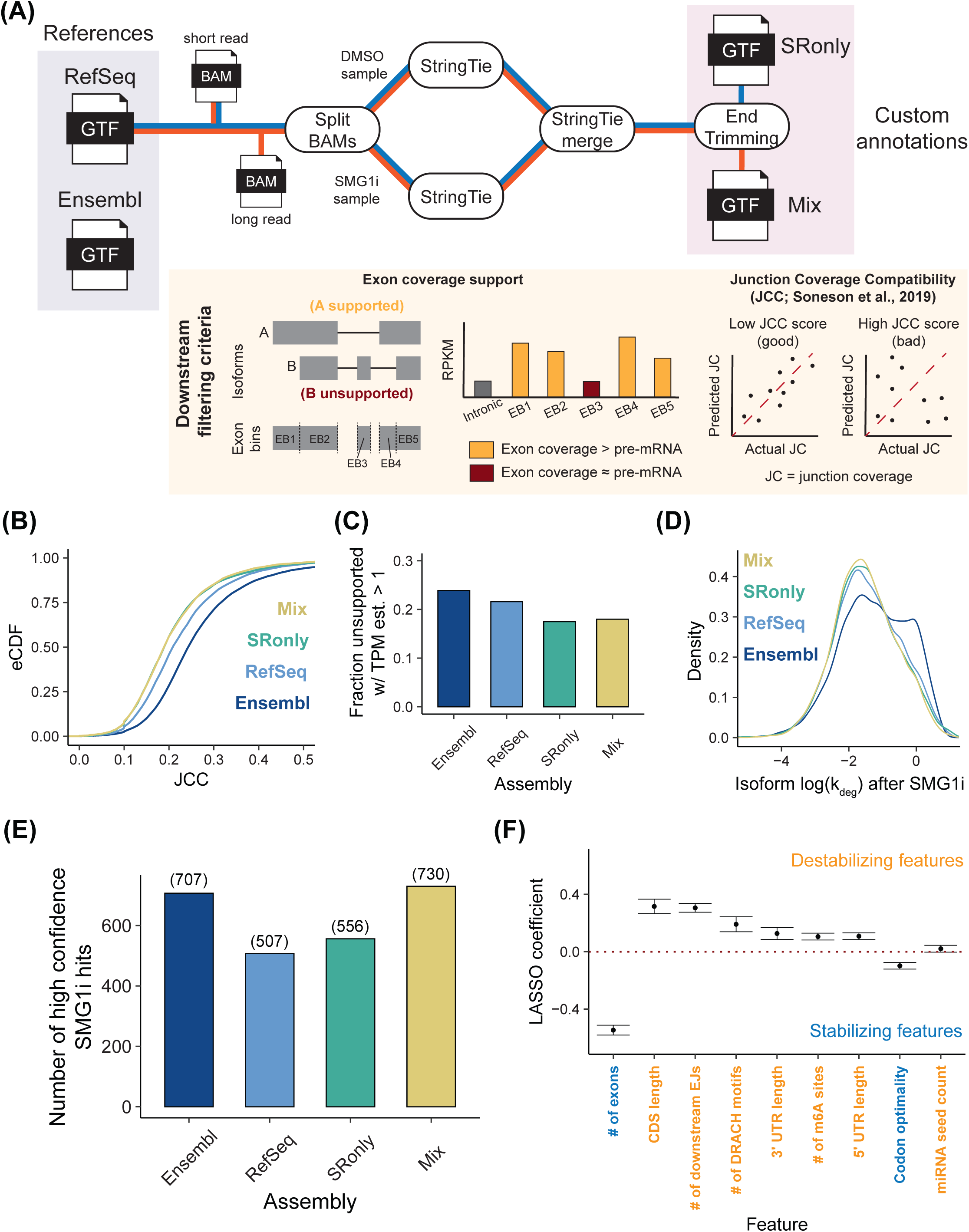
Annotation assembly and trimming improves isoform-level analyses. **(A)** Schematic of the AnnotationCleaner pipeline and downstream filtering criteria. Four annotations (2 references and 2 assemblies) were tested: Mix (StringTie using short and long read data), SRonly (StringTie using only short read data), RefSeq (hg38), and Ensembl (GRCh38 version 113). **(B)** Comparison of JCC score distributions for all four annotations tested. Higher JCC scores represent worse isoform abundance vs. junction coverage discrepancy. **(C)** Comparison of fraction of isoforms deemed “unsupported” by AnnotationCleaner that have an RSEM estimated TPM > 1 in all samples. Unsupported isoforms are those with at least one exon lacking greater than pre-mRNA levels of coverage in all samples. **(D)** Comparison of the distribution of degradation rate constant estimates in SMG1i treated cells, for all four annotations tested. **(E)** Comparison of the total number of high confidence SMG1i sensitive isoforms for each annotation. High confidence SMG1i sensitive isoforms are those with a L2FC(k_deg_) < −1, FDR < 0.01, and that pass both the unsupported isoform and JCC filtering (latter means expressed from a gene with JCC < 0.4). **(F)** LASSO regression model fit using the top performing annotation (Mix). Features are colored by whether increases in the feature value are associated with increases (destabilizing) or decreases (stabilizing) in k_deg_.

### AnnotationCleaner improves isoform-level analyses

We used AnnotationCleaner to build two annotations, one using only short read RNA-seq data (SRonly) and the other using both short and long read RNA-seq data (Mix), which has been shown to improve *ab initio* assembly [35]. In both cases, we used the RefSeq annotation to guide this process and RNA-seq data from both DMSO and SMG1i treated cells (**Figure 3A**). We then analyzed our NR-seq data with these two annotations and references from RefSeq and Ensembl [38,39]. Comparison of JCC score distributions revealed that AnnotationCleaner lowered average JCC scores, meaning that it improved the consistency between junction level coverage and isoform quantification (**Figure 3B**).

We then assessed the fraction of isoforms in each annotation with an RSEM-estimated TPM greater than one that had at least one exon with read coverage at or below that of pre-mRNA produced from the same gene. We term these “unsupported isoforms” and suspect that many of these are unexpressed isoforms erroneously assigned read coverage due to annotation discrepancies. Among the references, Ensembl had a higher fraction of such isoforms, which we believe is due to its higher gene model complexity and total isoform count (**Figure 3C** and **S3**). Both of the custom annotations had significantly lower rates of unsupported isoforms, suggesting that StringTie’s filtering and refinement of the annotation improved isoform quantification (**Figure 3C**).

To test whether annotation choice affects degradation rate constant inference, we assessed isoform turnover kinetics upon SMG1i treatment (**Figure 3D**). Mix, SRonly, and RefSeq annotations yielded similar degradation rate constant distributions, but the Ensembl analysis differed substantially. Specifically, the Ensembl analysis was plagued by a large number of apparently highly unstable isoforms that likely represent pre-mRNA (**Figure 3D, S3**, and **S4B**, left and middle panels). These spurious annotation-dependent results further support the efficacy of StringTie-based annotation filtering prior to isoform-level analyses of RNA-seq and NR-seq data. While the more conservative RefSeq annotation yielded better metrics than Ensembl, it also had far fewer high confidence NMD targets than either of the AnnotationCleaner annotations (**Figure 3E**). This highlights the benefits of expanding a more conservative reference with *ab initio* assembly. In addition, despite Ensembl having far more isoforms, it still yielded fewer high confidence NMD targets than the Mix assembly (**Figure 3E**). Finally, we note that while including long read data considerably improves identification of high confidence NMD targets, SRonly also had improved metrics relative to either of the references (**Figure 3C and D**). Thus, even if long read data is not readily available, short-read *ab initio* assembly can still significantly improve isoform quantification accuracy. All these observations led us to conclude that AnnotationCleaner improves EZbakR’s isoform-level NR-seq analyses.

Finally, we sought to further validate the isoform kinetic parameter estimates obtained using the top-performing AnnotationCleaner assembly (Mix). Towards that end, we evaluated whether our analyses aligned with other studies with respect to which features affect isoform stability. To associate transcript features with stability in DMSO-treated cells, we fit a LASSO regression model using our isoform degradation rate constant estimates (**Figure 3F**). We considered the impact of a number of features previously identified to correlate with RNA stability (length, number of exons, m6A sites, etc.). We found the strongest covariate with stability to be the number of exons, with more exons being associated with greater stability. This supports recent work suggesting that EJCs inhibit m6A deposition and stabilize isoforms [40–42]. In addition, we corroborated previous observations that overall transcript length and m6A levels negatively correlate with stability, and that codon optimality positively correlates with isoform stability [43]. We were also able to orthogonally validate these conclusions using previously published RNA-seq transcription inhibition data, suggesting that none of these correlations arise as an artifact of metabolic labeling (**Figure S4C** and **S4D**) [44]. Together, these analyses support the conclusion that isoform-level NR-seq analyses using our custom annotation accurately quantify isoform turnover kinetics.

### NMD is a major determinant of isoform decay rate variability within genes

NMD plays a key role in regulating isoform stability by selectively targeting PTC-containing and select non-PTC transcripts, where PTC transcripts are defined as those in which the termination codon (TC) occurs more than 50 nt upstream of the last exon junction [5]. However, the isoform-level kinetics of NMD have not been systematically measured, and how NMD fits into the overall kinetic landscape of mRNA degradation is unknown. To investigate these questions, we first compared the turnover kinetics of PTC and non-PTC isoforms under DMSO treatment (**Figure 4A**). Isoform-level degradation rate constants generally correlated with gene-level averages, but a subset of both PTC and non-PTC transcripts exhibited lower stability than their gene-level averages (higher isoform-level DMSO log(k_deg_) values), consistent with the characteristic instability of NMD substrates. The stability profiles of PTC and non-PTC isoforms were distinct (**Figure 4A**, median lines), with PTC isoforms often exhibiting much lower stabilities, as exemplified by SRSF3. These findings further highlight that gene-level averaging of stability measurements obscures isoform-specific variation in decay rate measurements across the transcriptome and that our analyses capture isoforms with distinct stabilities.

**Figure 4.**
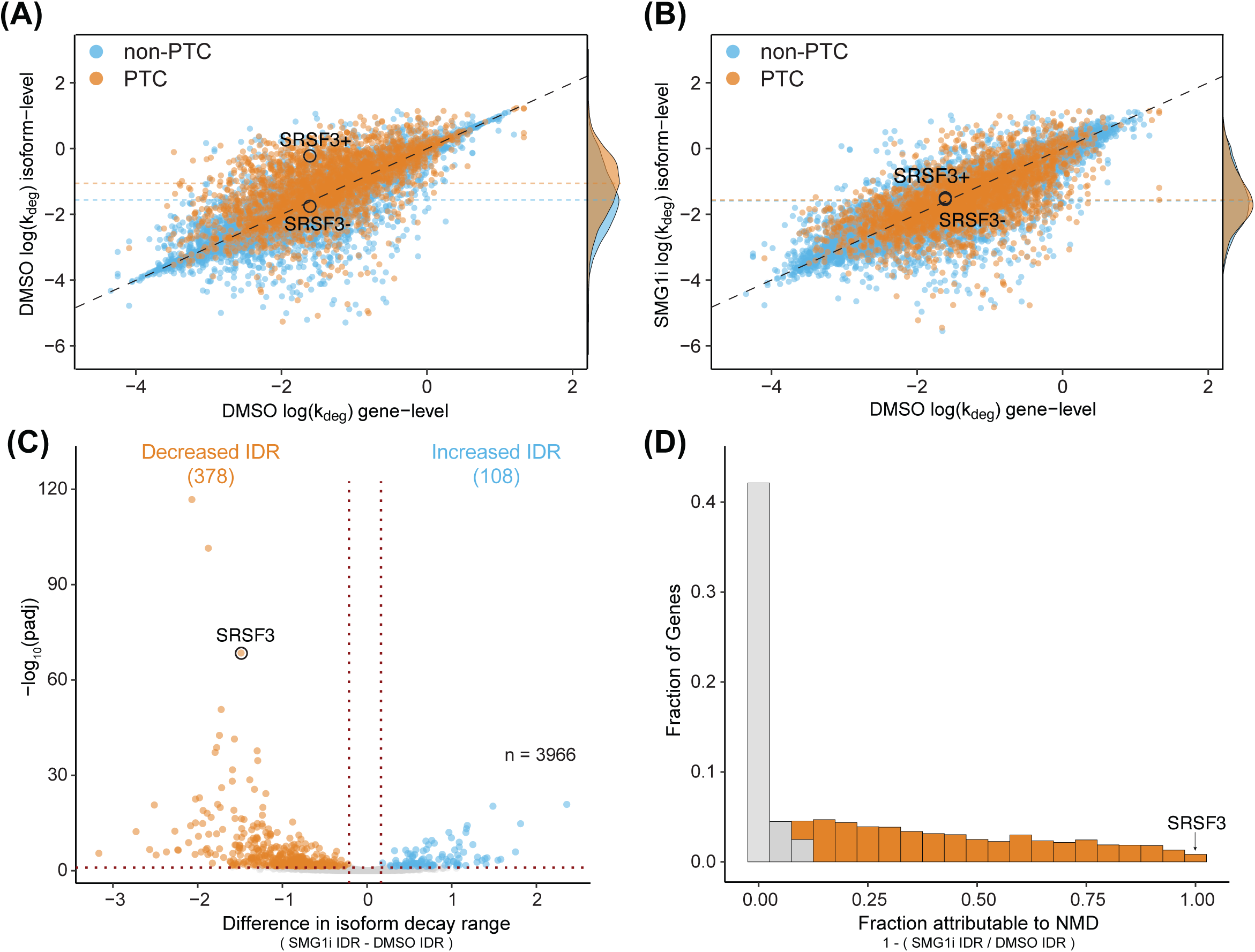
NMD plays a crucial role in driving variation in isoform stability within genes. (**A-B**) Scatter plots comparing isoform-level degradation rate constants (k_deg_) to gene-level degradation rate constants under DMSO (**A**) or SMG1i (**B**) treatment conditions. Isoforms are color-coded as non-PTC (blue) or PTC-containing (orange). The diagonal black line represents a linear relationship between gene-level and isoform-level k_deg_. Two isoforms of the SRSF3 gene are highlighted: the non-PTC isoform (SRSF3-) and the PTC-containing isoform (SRSF3+). Density plots on right y-axes show the distribution of PTC and non-PTC isoforms. Horizontal orange and blue lines indicate the median value for PTC and non-PTC isoforms. (**C**) Volcano plot showing the difference in isoform decay range (IDR) on the x-axis and −log_10_(padj) significance on the y-axis. Points represent multi-isoform genes, with significant differences defined by padj < 0.05 and IDR > 0.1 (blue) or IDR < 0.1 (orange). SRSF3 is highlighted in the text. (**D**) Histogram depicting the distribution of isoform stability differences attributable to NMD. Orange bars indicate cases where more than 10% of the stability difference is attributed to NMD. SRSF3 is discussed in the text.

We investigated the extent to which NMD explained instances of isoform-to-isoform stability variance within the same gene (**Figure 4B**). Following SMG1 inhibition, many isoforms were significantly stabilized, with their decay rate constants equal to or less than their gene-level averages (shift toward or below the diagonal). SMG1i treatment globally increased the stability of PTC isoforms, bringing their distribution of stabilities to match that of non-PTC isoforms (**Figure 4B**, right; compare PTC and non-PTC median lines). This indicates complete inhibition of NMD upon SMG1i treatment for many mRNAs. Given the prevalence of SMG1i-sensitive isoforms, we sought to directly quantify NMD’s transcriptome-wide contribution to RNA decay. To do this, we calculated an Isoform Decay Range (IDR) for each gene (see Methods for details), defined as the difference between the most and least stable isoforms within a gene. We then compared IDR changes between DMSO and SMG1i treatments (**Figure 4C**). This analysis revealed that of all multi-isoform genes, 378 genes exhibited a significant decrease in IDR, whereas only 108 showed an increase. To further estimate the impact of NMD on gene-level isoform stability differences, we calculated the fraction of gene-level IDR attributable to NMD (**Figure 4D**). This analysis revealed that 50.9% of genes with multiple well-expressed isoforms exhibited a decrease in IDR of 10% or more, with this difference in isoform stabilities attributable to SMG1i treatment. For instance, in SRSF3, nearly all stability differences between the PTC (SRSF3+) and non-PTC (SRSF3−) isoforms were driven by NMD, with SMG1i treatment increasing the stability of the PTC isoform to match that of the non-PTC isoform (**Figure 4D** and **4B**). More broadly, this analysis indicates that NMD influences isoform stability differences in over half of all multi-isoform genes expressed in HEK293 cells.

### Basal transcript instability constrains NMD

Leveraging our improved isoform-level measurements of NMD kinetics, we evaluated previously proposed rules for NMD. We separated transcripts into PTC and non-PTC isoforms and used linear regression to assess correlations between NMD efficiency and various parameters reported to influence NMD, including length of the coding sequence (CDS), TC-containing exon, longest internal exon, and 3’UTR. In doing so, we sought to determine whether the correlations observed in previous studies using RNA abundance data were evident using our measurements of turnover kinetics.

We found that PTC-containing isoforms with longer CDS lengths showed greater stabilization under SMG1i treatment compared to those with shorter CDS lengths, while non-PTC isoforms did not exhibit this correlation (**Figure S5A**). This supports previous findings that mRNAs with CDS lengths shorter than 100 nucleotides are less efficient NMD substrates [8], due to re-initiation at downstream start codons or other factors [45–47]. Also consistent with previous analyses, PTC-containing isoforms with more downstream exon-junctions (**Figure S5B**) and shorter distances between the termination codon and the nearest downstream exon-junction (**Figure S5C**) were more strongly stabilized by SMG1i [8,48].

We then examined the relationship between the length of TC-containing exons and changes in isoform stability upon NMD inhibition. Isoforms with PTCs in longer exons were associated with reduced sensitivity to SMG1i, which corroborates previous findings [8] (**Figure S5D**). Importantly, we also observed significantly lower basal stability of PTC isoforms with long TC-containing exons than those with short TC-containing exons (**Figure 5A**, left). This result suggests that transcripts with long TC-containing exons are subject to efficient NMD-independent turnover, to the extent that PTC-free transcripts with TC exon lengths greater than 1,500 nucleotides are often as unstable as PTC transcripts (**Figure 5A**, right). We further tested this idea by examining the correlation between basal transcript stability and the length of the longest internal exon (**Figure 5B**, left), finding a correlation for both PTC and non-PTC transcripts. Additionally, non-PTC isoforms with internal exons longer than 500 nucleotides show reduced basal stability similar to that of PTC-containing isoforms (**Figure 5B**, right). Together, our data suggest that the observed reduction in NMD susceptibility of transcripts with PTCs in long exons may be due to the destabilizing effects of long exons rather than ineffective NMD *per se*. The same principle can be applied to the related parameter of distance between the TC and the nearest downstream exon junction.

**Figure 5.**
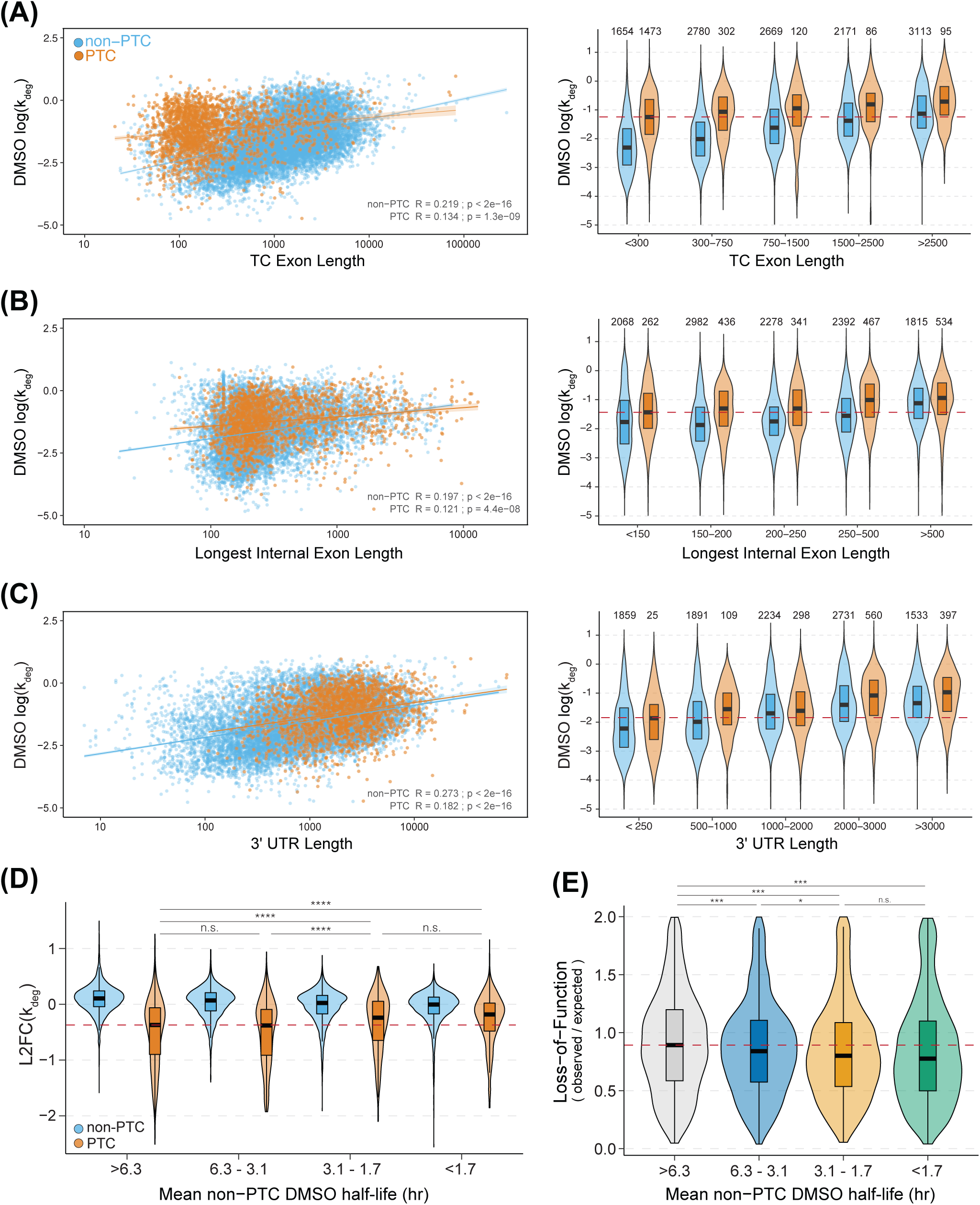
Intrinsic transcript instability limits NMD efficiency. (**A**-**C**) **Left:** Scatter plots depicting basal DMSO log(k_deg_) estimates as a function of (**A**) TC exon length, (**B**) longest internal exon length, and (**C**) 3’UTR length for non-PTC (blue) and PTC-containing (orange) isoforms. Linear regression fit lines with standard error shading are shown for both PTC and non-PTC data, along with corresponding correlation coefficients (R) and significance values (p-value) for each regression. **Right:** Violin plots displaying the distribution of basal DMSO log(k_deg_) estimates against length-wise binning of the same features (**A**-**C**) for non-PTC (blue) and PTC-containing (orange) isoforms. Dashed red lines denote the mean basal DMSO log(k_deg_) of the shortest feature length of PTC isoforms. (**D**) Violin plots showing the distribution of L2FC(k_deg_)’s between SMG1i and DMSO treatments for non-PTC (blue) and PTC-containing (orange) isoforms. Isoforms are binned based on the mean DMSO log(k_deg_) of non-PTC isoforms. The horizontal red dashed line represents the lowest median L2FC(k_deg_) observed across all bins. (**E**) Violin plots displaying the distribution of loss-of-function (observed/expected) scores for non-PTC isoforms, binned according to the mean DMSO log(k_deg_) of non-PTC isoforms. Statistical significance of pairwise comparisons between bins is indicated above the plots (n.s. not significant, * p < 0.05, ** p < 0.01, *** p < 0.001).

Lastly, we analyzed 3’UTR length, a common characteristic implicated in decay of EJC-independent NMD substrates. In our analysis, we observed a slight inverse correlation between 3’UTR length and the response of PTC or non-PTC isoforms to SMG1i (**Figure S5F**). However, when comparing 3’UTR length with basal RNA stability, we found a much stronger correlation, where longer 3’UTRs were associated with reduced basal RNA stability (**Figure 5C**, left). Both PTC-containing and non-PTC isoforms followed this trend, with non-PTC isoforms with 3’UTRs longer than 2,000 nucleotides showing destabilization comparable to most PTC isoforms (**Figure 5C**, right). These results suggest that isoforms with longer exons (TC-containing and internal) and 3’UTRs are prone to decay by other pathways, potentially reducing their sensitivity to NMD.

To globally evaluate how basal stability constrains NMD, we took advantage of the ability to measure the decay rate constants of PTC and non-PTC isoforms produced from the same gene. We binned genes based on the mean DMSO half-lives of their non-PTC isoforms and compared decay rate constant changes between SMG1i and DMSO treatments for both PTC and non-PTC isoforms (**Figure 5D**). PTC-containing isoforms showed progressively less responsiveness to SMG1i as the basal stability of non-PTC isoforms from the same gene decreased (**Figure 5D**). One example of basal instability influencing NMD sensitivity is HMGXB4. In this gene, the inclusion of a poison exon (i.e. an exon that encodes a PTC when included in mature mRNA), along with an upstream exon, creates an open reading frame that introduces a stop codon within the poison exon (**Figure S5G**). We found that all isoforms of HMGXB4 are highly unstable (median isoform half-life of 53 mins), with both non-PTC and PTC-containing isoforms exhibiting similar expression levels and higher decay rate estimates under DMSO treatment (**Figure S5G**, right; high ratio of s^4^U labeled to unlabeled reads). Furthermore, SMG1i treatment did not stabilize any of the HMGXB4 isoforms, including the poison exon-containing isoform (**Figure S5G**, right). Taken together, these findings underscore the potential for competing RNA decay pathways to limit the capacity of NMD to regulate gene expression.

We next asked whether quality control may also be impaired by basal transcript instability, as previously suggested [8]. We used the gnomAD database to obtain gene-level loss of function observed/expected upper bound fraction (LOEUF) scores [49], a measure of a gene’s tolerance to inactivating mutations. We binned each gene by its mean non-PTC isoform DMSO half-life (**Figure 5E**) and found decreased LOEUF scores among genes with shorter mean non-PTC isoform DMSO half-lives. The finding that genes with lower basal stability tend to be less tolerant to loss-of-function mutations is predicted by a model in which NMD is unable to effectively reduce the abundance and translation of PTC-containing transcripts from genes with low basal transcript stability.

### NMD efficiently targets a subset of non-PTC isoforms, including a novel class of PTC-free AS-NMD substrates

Current dogma suggests that decay of PTC-containing transcripts is inherently more efficient than decay of non-PTC NMD targets, due to decay stimulation by the EJC. By utilizing degradation rate constant measurements, rather than solely relying on RNA abundance changes, we could better evaluate the differences in stabilities between PTC and non-PTC NMD substrates. To make this comparison, we curated two sets of previously validated PTC-dependent and -independent NMD substrates. The first set consisted of transcripts containing previously identified conserved poison exons [50]. The second set included genes encoding NMD factors, most of which are autoregulated by the pathway, along with the previously characterized NMD substrate IRE1 (ERN1). Several of these genes have 3’UTRs that have been shown in reporter assays to be sufficient to trigger decay in the absence of a PTC [51–54]. To identify additional high-confidence, SMG6-dependent NMD targets transcriptome-wide, we performed Akron5-seq [55] to quantitatively capture RNA 5’ ends at single-nucleotide resolution. To stabilize decay intermediates, the major 5’-3’ exonuclease XRN1 was depleted alone or in combination with SMG6. Our analysis revealed that SMG6-dependent peaks were enriched at both PTCs and regulated non-PTC termination codons and that decapping increased upon SMG6 depletion (**Figure S6A**), consistent with previous SMG6-cleavage mapping studies [56–58]. We compared changes in peak abundance between XRN1-depleted and XRN1/SMG6 co-depleted conditions with changes in k_deg_ following SMG1 inhibition (**Figure S6B**). We also analyzed a published dataset of SMG6 cleavage sites to further corroborate our findings [56]. This analysis enabled us to find a stringently filtered set of non-PTC isoforms that depend on SMG6 cleavage near termination codons, adhere to the 50-nucleotide rule, and exhibit sensitivity to SMG1 inhibition (**Figure S6C-F**; see Methods for details).

Next, we compared the basal stabilities and changes in decay rate constants upon SMG1i treatment for all predicted PTC and non-PTC isoforms (**Figure 6A**). Globally, PTC-containing transcripts were generally more unstable and more strongly stabilized by SMG1i than non-PTC isoforms. Overlaying the aforementioned set of PTC and non-PTC decay substrates revealed that the set of poison exon-containing isoforms was, as expected, on average highly unstable under DMSO conditions and strongly stabilized by SMG1i. Notably, a subset of non-PTC NMD isoforms, including from the TSC22D3, PDRG1, NUDT16L1, CDKN2AIP, and RASSF1 genes, were at least as unstable as validated PTC-containing isoforms and were similarly strongly stabilized by SMG1i. These findings suggest that NMD can be just as effective in targeting certain non-PTC substrates as it is for PTC isoforms.

**Figure 6.**
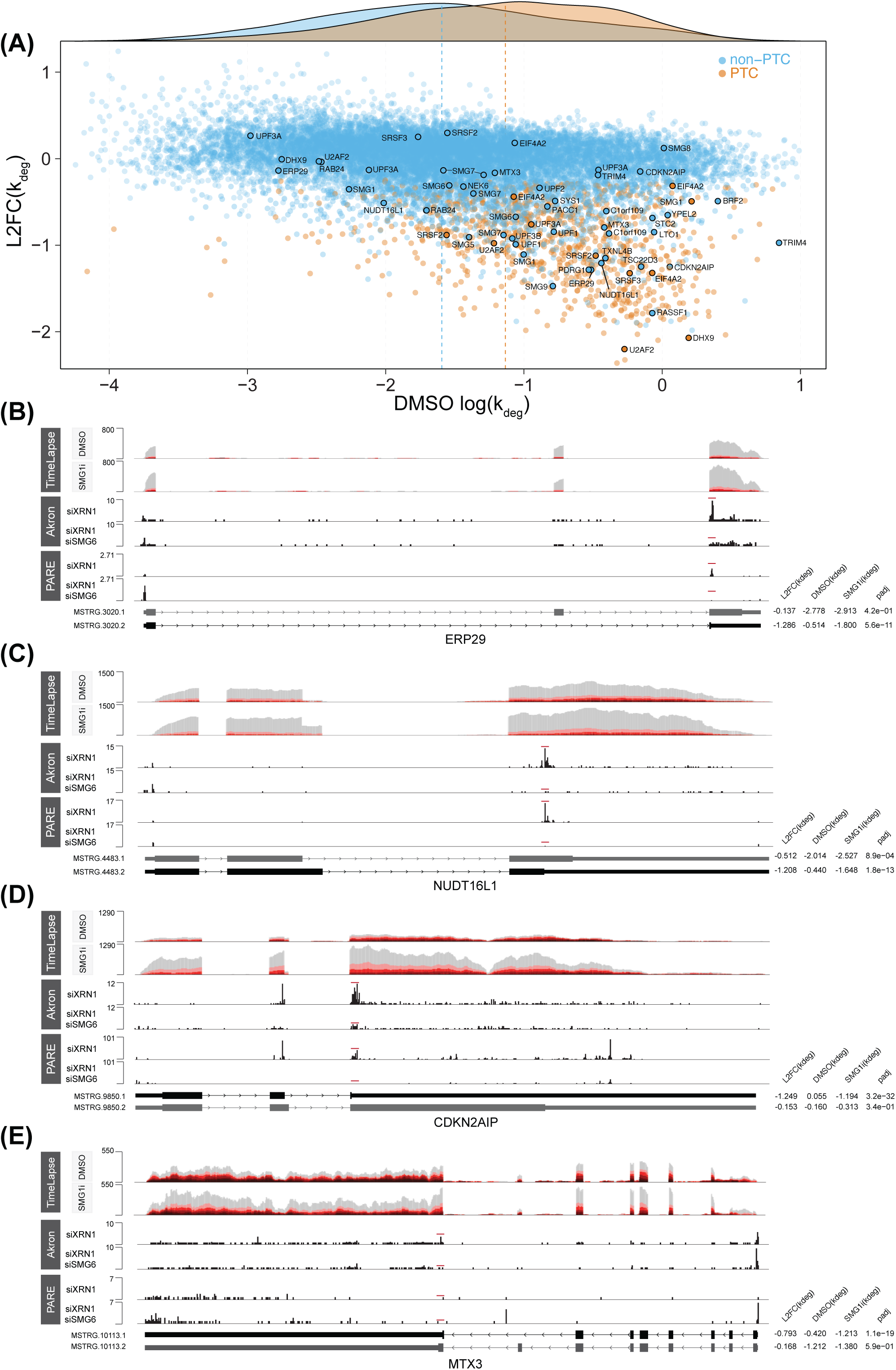
NMD efficiently regulates a subset of EJC-independent substrates. (**A**) Scatter plot comparing the L2FC(k_deg_)’s between SMG1i and DMSO treatments (x-axis) to the basal DMSO log(k_deg_) estimates (y-axis) for non-PTC (blue) and PTC-containing (orange) isoforms. *Top*: Density plot displaying the distribution of DMSO log(k_deg_) estimates for PTC and non-PTC isoforms, with vertical lines indicating the median of each group’s distribution. Genes highlighted in the figure are discussed further in the text and in Figure S6. (**B**-**E**) **Top:** TimeLapse-seq tracks for ERP29 (**B**), NUDT16L1 (**C**), CDKN2AIP (**D**), and MTX3 (**E**) under DMSO (upper tracks) and SMG1i (lower tracks) treatment conditions. The degree of s^4^U labeling is indicated by increasing darkness of red, corresponding to the number of T-to-C mutations per read, with unlabeled reads shown in gray. **Bottom:** Akron-seq (this study) and PARE-seq [56] tracks from XRN1 and XRN1+SMG6 co-depletion experiments, marking SMG6-dependent cleavage sites in Akron-seq by red lines above the sequence tracks. The isoform from each gene is schematized below, with the ORFs represented by thicker lines. SMG1i-sensitive isoforms are colored black, while - insensitive isoforms in gray. Corresponding decay rate measurements for these isoforms are displayed to the right.

To orthogonally validate our findings, we assessed transcript stability by exclusively analyzing reads mapping to splice junctions, an approach that is less dependent on annotation quality and avoids potential rate constant biases from high turnover pre-mRNA. For this, we leveraged the ability of the EZbakR-suite to flexibly analyze reads mapping to various transcript features, including reads that span exon-exon splice junctions. Junction-level and isoform-level measurements revealed highly similar patterns of PTC and non-PTC transcript degradation by NMD, further supporting the accuracy of our isoform-level stability estimates **(Figure S6G-J)**.

Isoform- and junction-level analyses consistently indicated that mRNAs with conserved poison exons have a broad range of half-lives in unperturbed cells, spanning from 45 mins to over two hours. Correspondingly, the curated set of PTC isoforms also responded variably to SMG1i treatment. These findings reinforce the conclusion that there are substantial differences in NMD efficiency even among high-confidence PTC transcripts. Junction-level analyses also supported the existence of highly efficient NMD of certain non-PTC transcripts, as nearly all curated non-PTC target mRNAs exhibited basal stabilities and responses to SMG1i within the interquartile ranges of the curated set of PTC mRNAs in both isoform- and junction-level analyses.

These analyses also revealed unexpected examples of genes for which differential termination codon positioning in terminal exons drives PTC-free alternative splicing-linked NMD (AS-NMD) (**Figure 6B-E**). ERP29, NUDT16L1, CDKN2AIP, and MTX3 each are alternatively spliced to express two isoforms, one SMG1i sensitive and one insensitive (**Figure S6H**). NMD-sensitive isoforms of these genes contain shortened open reading frames and extended 3’UTRs, creating TCs that are efficiently recognized by NMD despite adhering to the 50-nucleotide rule. This demonstrates how cells can achieve robust isoform-specific regulation by coupling alternative splicing to efficient NMD in a downstream exon junction-independent manner.

### Machine learning identifies covariates of isoform-specific NMD kinetics

To identify covariates of PTC-dependent and independent NMD, we identified a subset of transcripts that were either high-confidence instances of SMG1i sensitivity or insensitivity and then fit a LASSO binary classification model to either the set of all PTC or non-PTC containing isoforms of these subsets. The model included a mix of established factors from the literature and new variables that we hypothesized could influence NMD (**Table S5**).

As expected, the strongest predictor of PTC-containing isoform NMD efficiency was the presence of a downstream exon junction (**Figure 7A**). The model also predicted other important factors previously linked to decay of PTC-containing RNAs [8,10], including the number of downstream exon junctions, the distance between the PTC and the mRNA 5’ end, and 3’UTR length, confirming our approach to transcript classification and machine learning. However, features recently proposed to be positively associated with NMD susceptibility due to pervasive mis-splicing, such as total exon number and intron length, did not significantly contribute to our model’s predictions [59]. CDS length had opposite correlations in PTC and non-PTC isoforms: longer CDS lengths were positively correlated with NMD sensitivity in PTC-containing isoforms, whereas shorter CDS lengths correlated with lower NMD efficiency in non-PTC isoforms (**Figure 7A** and **7B**). This demonstrates that our transcript classification can effectively distinguish between correlates of PTC and non-PTC NMD substrates, capturing both shared and distinct regulatory features.

**Figure 7.**
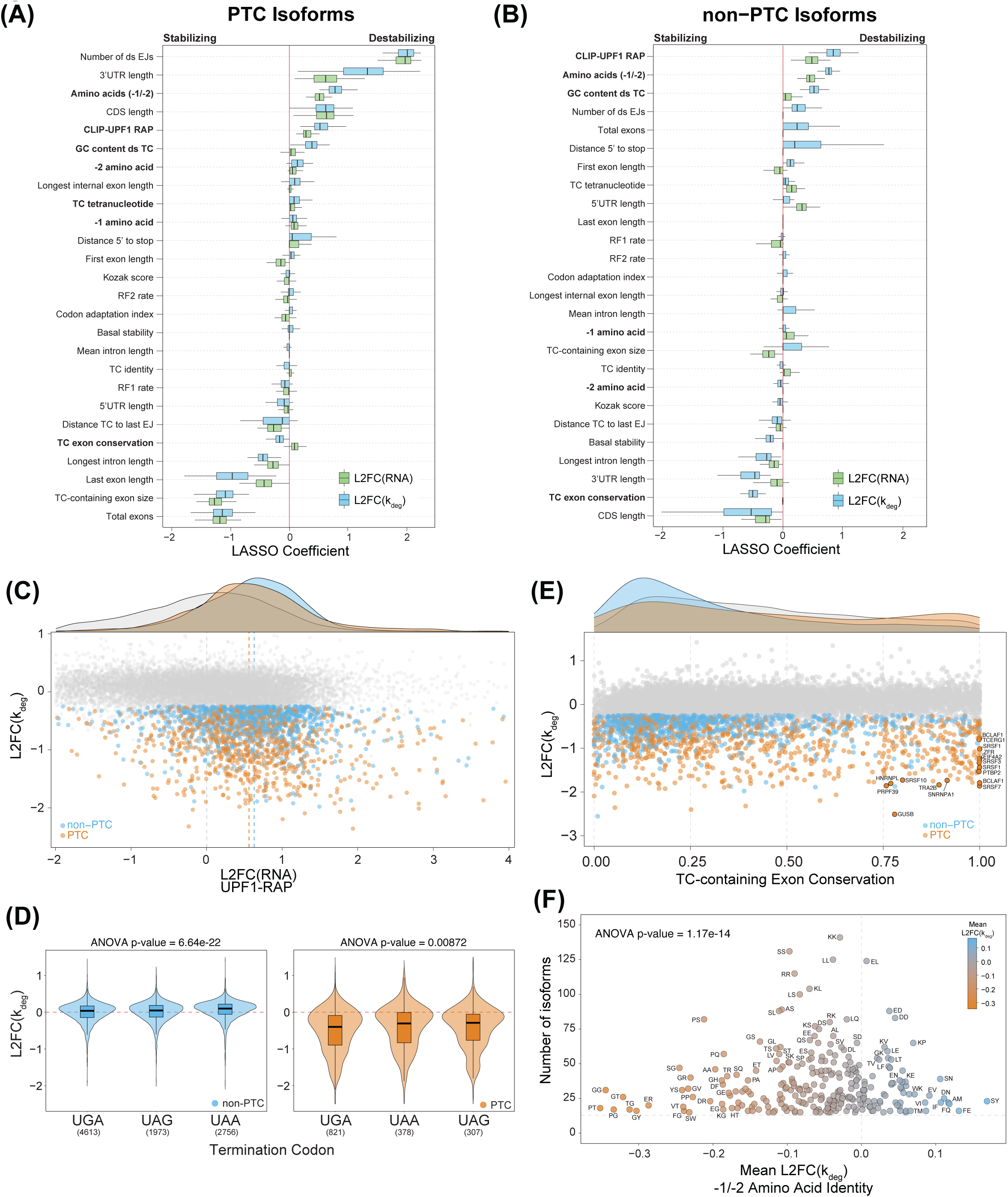
Machine learning reveals key features of PTC and non-PTC isoform-specific NMD kinetics. (**A-B**) Box plots showing the estimated LASSO regression coefficients (100 bootstrapped resamplings) for L2FC(k_deg_) (blue) and L2FC(RNA) in PTC (**A**) and non-PTC (**B**) isoforms for tested features (left). (**C**) Scatter plot showing L2FC(k_deg_) between SMG1i and DMSO treatments versus L2FC(RNA) enrichment from CLIP-UPF1 RNA affinity purification (RAP). Non-PTC isoforms are represented in blue, PTC isoforms in orange, and non-responsive isoforms in gray. Vertical lines indicate the median of each group’s distribution. **Top:** Density plot of each group with vertical lines indicating the median of each group’s distribution. (**D**) Violin plots showing the distribution of L2FC(k_deg_) between SMG1i and DMSO treatments for different TC identities. Left plots (blue) are for non-PTC isoforms and right plots (orange) are for PTC isoforms. The horizontal dashed red line represents no change in the k_deg_ between conditions. Number of observations for each stop codon (TC) identity are shown below the plots. ANOVA p-value indicates statistical significance after assessing whether the variance between TC groups exceeds the variance within TC groups (i.e., whether there is strong evidence that different TC groups have different average L2FC(k_deg_)’s). (**E**) Scatter plot of L2FC(k_deg_) versus TC-containing exon conservation scores. Non-PTC isoforms are shown in blue, PTC isoforms in orange, and non-responsive isoforms in gray. **Top:** Density plot for each group, with vertical lines indicating the median of each distribution. A subset of highly conserved TC-containing exon isoforms is labeled. (**F**) Scatter plot of mean L2FC(k_deg_) for each C-terminal amino acid pair versus the number of isoforms containing them. Amino acid pairs are colored by their mean L2FC(k_deg_), as indicated in the legend (*top right*). ANOVA p-value reflects statistical significance as described in (D).

### UPF1 association correlates with decay of PTC and non-PTC transcripts

To further assess how various features relate to NMD susceptibility of PTC- and non-PTC-containing isoforms, we examined RNA-protein interactions using our previously published UPF1 RNA affinity purification and sequencing (RAP-seq) dataset [60,61]. We observed a strong positive correlation between UPF1 recovery and the decay of both PTC- and non-PTC-containing transcripts (**Figure 7C**). Notably, previously characterized PTC and non-PTC NMD substrates were enriched in UPF1 RAP-seq data (**Figure S7A**), supporting prior findings that UPF1 preferentially associates with NMD-susceptible mRNAs [62,63].

### Nucleotide and amino acid sequences associated with efficient NMD

In addition to RNA-protein interactions, we identified multiple mRNA sequence-based features that influence NMD susceptibility. We observed a significant difference in GC content within the 200 nucleotides downstream of termination codons, which was strongly associated with stability changes but not RNA abundance changes (**Figure 7A** and **7B**). Non-responsive isoforms exhibited a median GC content of 44%, whereas responsive PTC and non-PTC isoforms had significantly higher GC contents at 50% and 54%, respectively (**Figure S7B**), consistent with previous findings on the role of 3’UTR GC content in modulating NMD efficiency [64,65].

Termination codon identity was significantly associated with differential SMG1i sensitivity, with both PTC and non-PTC isoforms containing UGA stop codons exhibiting the strongest response to SMG1i treatment (**Figure 7D**). Further, LASSO regression identified TC tetranucleotide context (TC identity plus the following +4 nucleotide) as a stronger correlate of PTC and non-PTC NMD efficiency than TC identity alone. Among non-PTC targets, mRNAs with UGAC tetranucleotides were most strongly affected by SMG1i treatment. Our evidence of preferential decay of endogenous transcripts with UGA TCs, particularly those with a +4 C nucleotide, corroborates the results of a recent massively parallel reporter assay of EJC-dependent and independent substrates [66] (**Figure S7C**). Notably, UGAC stop codons support less efficient termination than other tetranucleotide contexts, suggesting that the NMD machinery may be better able to recognize termination events with slow kinetics [67–70].

The identification of several sequence features associated with NMD susceptibility suggested that mRNA sequences may evolve to tune NMD efficiency. In support of this hypothesis, we found that sequence conservation of the exon containing the stop codon was strongly anti-correlated with RNA stability in both PTC and non-PTC isoforms, whereas RNA abundance showed little to no enrichment for exon conservation scores. We observed a bimodal distribution of SMG1i-sensitive PTC isoforms, consisting of a large population of poorly conserved exons and a notable subset of very highly conserved exons, many corresponding to known poison cassette exons in RNA-binding protein genes (e.g. SR proteins, PTBP2, EIF4A2, and BCLAF1) (**Figure 7E**). These findings reflect the diverse ways in which human cells use NMD to both maintain gene expression fidelity and enforce gene expression regulation.

Finally, the model also identified C-terminal dipeptide sequence context as a correlate of NMD susceptibility (**Figure 7A** and **7B**). Both PTC and non-PTC isoforms exhibited enrichment of specific residues at the minus one and minus two positions relative to the TC, with glycine, threonine, cysteine, and serine being enriched at the minus one position, while glycine, cysteine, and proline were enriched at the minus two position (**Figures S7D, S7E**). Notably, glycine and proline were particularly enriched in many of the dipeptide identities that were highly sensitive to SMG1i treatment (**Figure 7F**). This enrichment pattern resembles findings from a massively parallel reporter assay of EJC-independent substrates [66], further suggesting that amino acid sequence context plays a significant role in modulating transcript stability.

Taken together, these analyses highlight the power of kinetic measurements in resolving NMD covariates. Features such as stop codon exon conservation and downstream GC content, which were strong correlates in our kinetic data, were either weakly correlated or displayed opposite trends when analyzed using RNA abundance. These findings reinforce the importance of isoform-specific decay kinetics in understanding transcriptome-wide NMD regulation and reveal a complex interplay among RNA sequence features, exon conservation, and UPF1 association in shaping NMD efficiency across diverse transcript classes.

## Discussion

RNA abundance measurements have been instrumental in shaping our understanding of gene expression mechanisms. However, RNA abundance is an indirect readout and can generate an incomplete or misleading picture of the contributions of RNA biogenesis and decay to gene expression regulation. In this work, we present the first strategy capable of inferring transcript isoform synthesis and degradation kinetics from short-read NR-seq of total RNA. We have implemented this strategy in our recently developed EZbakR-suite to facilitate its wide adoption among researchers. We envision that NR-seq and EZbakR will be powerful and broadly applicable tools to understand mechanisms and cellular functions of gene expression regulatory pathways.

Our findings underscore how the choice of annotation can significantly impact isoform-level investigations of RNA-seq data. We have developed AnnotationCleaner to increase representation of PTC isoforms that are often omitted from reference annotations while culling spurious transcript models. We find that it is particularly important to limit the contribution of isoforms corresponding to unstable pre-mRNAs when estimating mature mRNA decay rate constants, an issue we find to be prevalent in Ensembl annotations. For most applications, we advocate for starting from a more conservative annotation and expanding it with tools like StringTie to generate custom references for each biological system of interest. In the Mix annotation used for this work, we have emphasized accuracy over completeness, but different experimental systems or research questions may require different approaches to annotation assembly and filtering.

While previous studies have primarily inferred NMD activity from changes in steady-state RNA levels, our findings emphasize the need for direct kinetic measurements to fully capture the dynamics of NMD-mediated transcript turnover. Using EZbakR, we analyzed transcript synthesis and degradation kinetics at isoform and sub-isoform resolution, integrating NR-seq with acute NMD inhibition to reveal the kinetic effects of NMD. Our findings reveal that while PTC-containing isoforms are generally susceptible to NMD, conserved poison exon-containing isoforms exhibit substantial variability in decay rates. Like other studies, we find that increased numbers of downstream exon junctions and increased 3’UTR length are associated with greater likelihood of efficient decay of PTC-containing transcripts. Importantly, we also identify a subset of non-PTC isoforms that are as unstable as PTC-containing transcripts and are equally stabilized upon SMG1i treatment, challenging the assumption that non-PTC isoforms are necessarily less efficiently targeted by NMD. These include what are to our knowledge the first known examples of PTC-free AS-NMD in mammalian cells. These insights expand our understanding of NMD-based regulation and highlight the complexity of transcript features that dictate susceptibility to degradation.

The potential for basal transcript instability (i.e., the efficiency with which non-NMD decay pathways act on a transcript) to constrain NMD has been previously recognized [8]. However, EZbakR offers the important advantage of providing accurate measurements of the stability of PTC and non-PTC isoforms from the same gene. This approach identifies PTC-containing transcripts that are likely resistant to NMD due to competing turnover mechanisms and reveals mechanisms contributing to known NMD efficiency correlates. Notably, our data suggest that inefficient decay of transcripts with long PTC-containing exons is largely attributable to basal instability. We find the mRNAs with long internal exons, irrespective of TC position, are often as unstable as NMD-sensitive PTC transcripts. In addition, our data provide a mechanistic explanation for recent observations that loss of function-intolerant genes are less sensitive to NMD [9,49].

Putting NMD in the context of global transcript decay rate constants also sheds light on the regulation of mRNAs with long 3’UTRs. We find that transcripts with long 3’UTRs, like those with long internal exons, often exhibit decay rate constants similar to those of well-targeted PTC transcripts. These results are consistent with a recent report of NMD-independent decay of long 3’UTRs [71]. Thus, effective 3’UTR-based NMD may rely not only on sensitivity to NMD but also insensitivity to other decay pathways. These observations, together with previous identification of RNA-binding proteins capable of preventing NMD of transcripts with long 3’UTRs [72,73], suggest a complex set of determinants for decay of transcripts with long 3’UTRs and highlight the importance of considering competing decay pathways when drawing mechanistic conclusions from transcriptome-wide RNA abundance correlations.

Our data show that NR-seq and isoform-level EZbakR analyses can reveal distinct kinetic signatures of various RNA decay pathways and explain how different turnover processes interact to control basal RNA stability. Looking ahead, EZbakR analyses of NR-seq data can be used to dissect the kinetic mechanisms of many gene expression regulatory processes. For example, integration of EZbakR with RNA-binding protein (RBP) perturbation studies can identify whether specific RBPs modulate splicing decisions, transcript stability, or both. Applying isoform-level NR-seq to diverse cell types and disease models will help distinguish tissue-specific regulation from pathological disruptions in RNA processing, providing insights into disease mechanisms. These advances in understanding isoform-specific degradation kinetics also have practical applications, as they promise to improve our understanding of the nucleotide sequence determinants of RNA stability, knowledge that could guide the rational design of stable therapeutic mRNAs for gene therapy and RNA-based medicines.

## Conclusions

In this work, we expand the EZbakR-suite to enable isoform-level kinetic inference from short-read NR-seq data and introduce a strategy for refining and filtering transcript annotations, which is critical for accurately quantifying kinetic rate constants. Using the EZbakR-suite and acute NMD inhibition, we generate the first isoform-resolution kinetic map of human NMD. Our analyses uncover unexpected variability in the NMD efficiency of conserved poison exon-containing transcripts, the efficient targeting of non-PTC isoforms by NMD, and the impact of competing RNA decay pathways on both the regulatory and quality control functions of NMD. This study illustrates how isoform-level analyses with the EZbakR-suite can deepen our understanding of RNA kinetic regulation and provides a framework for investigating transcript-specific degradation kinetics, paving the way for future studies on how decay pathways shape the human transcriptome.

## Methods

### Small-molecule SMG1i synthesis

Small-molecule SMG1 kinase inhibitor (11j) was synthesized as previously described [21].

### SMG1i-treatment and TimeLapse chemistry

Human HEK293 cells were cultured at 37°C under 5% carbon dioxide in a humidified chamber in Dulbecco’s modified Eagle medium (DMEM) supplemented with 10% v/v fetal bovine serum (FBS) and 1% v/v penicillin-streptomycin. Cells were seeded at 0.3 × 10⁶ cells per well in eight wells across two 6-well plates the day before treatment. Cells were treated with either 1 µM SMG1i (four wells) or DMSO (four wells) for eight hours. Additionally, 100 µM s^4^U (Cayman Chemical 16373) was added to three wells from each condition for the last two hours. Media was removed, and total RNA was extracted using TRIzol Reagent. The TimeLapse workflow was then followed as previously described [13,76].

### rRNA depletion for TimeLapse RNA-seq libraries

rRNA depletion of 1 µg total RNA was performed as previously described [77]. For library preparation, 100 ng of ribodepleted RNA was used as input for the NEBNext Ultra II Directional RNA Library Prep Kit (E7760). Libraries were prepared following the manufacturer’s instructions. Libraries were sequenced by Admera Health, on the Illumina NovaSeq X+ platform at a depth of 120M reads per sample.

### PacBio Full-Length Transcript Sequencing (Iso-Seq)

PacBio Iso-Seq was performed by CD Genomics. A total of 2 μg RNA was used as input for RNA sample preparation. First-strand cDNA synthesis, including reverse transcription and template switching, was carried out using the Clontech SMARTer™ PCR cDNA Synthesis Kit and the PacBio Iso-Seq Express Oligo Kit, followed by PCR amplification. SMRTbell libraries were generated using the SMRTbell Template Prep Kit. Sequencing was performed on a single sample of SMG1i and DMSO, with a total read depth of 20 million reads. RNA sequencing data processing was carried out by first converting the BAM files into FASTQ format using the bam2fastq [78] tool for both DMSO and SMG1i samples. The resulting FASTQ files were aligned to the hg38 reference genome using minimap2 [79] with the splice-aware option, producing SAM files [*minimap2 -ax splice -ub hg38.fa ccs.fastq.gz > ccs.sam*]. The resulting SAM files were then converted to sorted BAM files using SAMtools [80] [*samtools view -bS ccs.sam | samtools sort - o .ccs_sorted.bam*].

### AnnotationCleaner pipeline

AnnotationCleaner is a Snakemake pipeline that uses StringTie2 [35] in conjunction with custom scripting to build and refine custom annotations. It supports the use of short and long read RNA-seq data. StringTie2 is used to assemble annotations for each sample (or pair of related short- and long-read samples), as well as to merge the individual assemblies. The merged assembly is then split into exonic and intronic bins, and featureCounts [81] is used to quantify the coverage of each of these bins (sizes of these bins can be adjusted; default used in this paper was for exonic bins to be 200 nt in length and intronic bins to be 600 nt in length). Exonic bins cover regions that are exonic in at least one annotated isoform, and intronic bins cover regions that are intronic in all annotated isoforms.

The intronic coverage of each gene is computed (in units of RPKM), and noisy estimates are regularized using genome-wide trends. More specifically, a regression model of exonic coverage vs. intronic coverage is fit, and the trendline is used as an informative prior to regularize gene-specific estimates. This specific strategy is identical to the replicate variability regularization strategy implemented in bakR [82] and EZbakR [17].

The coverage of each exonic bin is compared to the intronic coverage via a negative binomial null hypothesis test, and the resulting p-value is multiple test adjusted using the method of Benjamini and Hochberg. Ends are trimmed by removing terminal exonic bins deemed poorly supported, and isoforms with unsupported exonic bins whose removal would introduce new splice junctions are left untrimmed and flagged in the final GTF file output by AnnotationCleaner (boolean column added called “problematic”).

### Isoform CDS and PTC annotations

The factR2 [83] package was used to characterize alternative splicing events, annotate CDSs, and predict isoforms containing PTCs. A factRObject was created from the transcript annotation file (SR-only.gtf or Mix.gtf), using the human genome as the reference. Annotated Ensembl CDSs were matched to transcripts in the custom annotation. The factR2-processed annotation was then passed to SQANTI3 (v5.2.2) [84] for further identification of novel CDSs using the GeneMarkS-T (GMST) [85] algorithm, using GENCODE v37 as the reference annotation and hg38 as the genome reference. To ensure accuracy, gffread [86] was used to correct any GTF anomalies, and Orfanage [87] was employed to integrate annotated ORFs from factR2 and SQANTI3. Finally, the refined annotation was reprocessed through factR2 to identify additional PTC-containing isoforms from all factR2 and SQANTI3 identified CDSs.

### Transcript isoform kinetic parameter estimation

We have implemented a novel linear mixing model in EZbakR, designed to estimate isoform-specific kinetic parameter estimates. It accepts as input transcript isoform abundance estimates and fraction new estimates for each transcript equivalence class (TEC). The former can come from any isoform quantification tool (RSEM is implemented in fastq2EZbakR and was used in all analyses presented here), and the latter is most easily obtained from EZbakR’s EstimateFractions() function, used in this work (default settings except pold_from_nolabel was set to TRUE, features was set to c(“XF”, “TEC”), and filter_condition was set to ‘|’). For each gene, the EstimateIsoformFractions() function fits the following linear mixing model using the method of maximum likelihood:

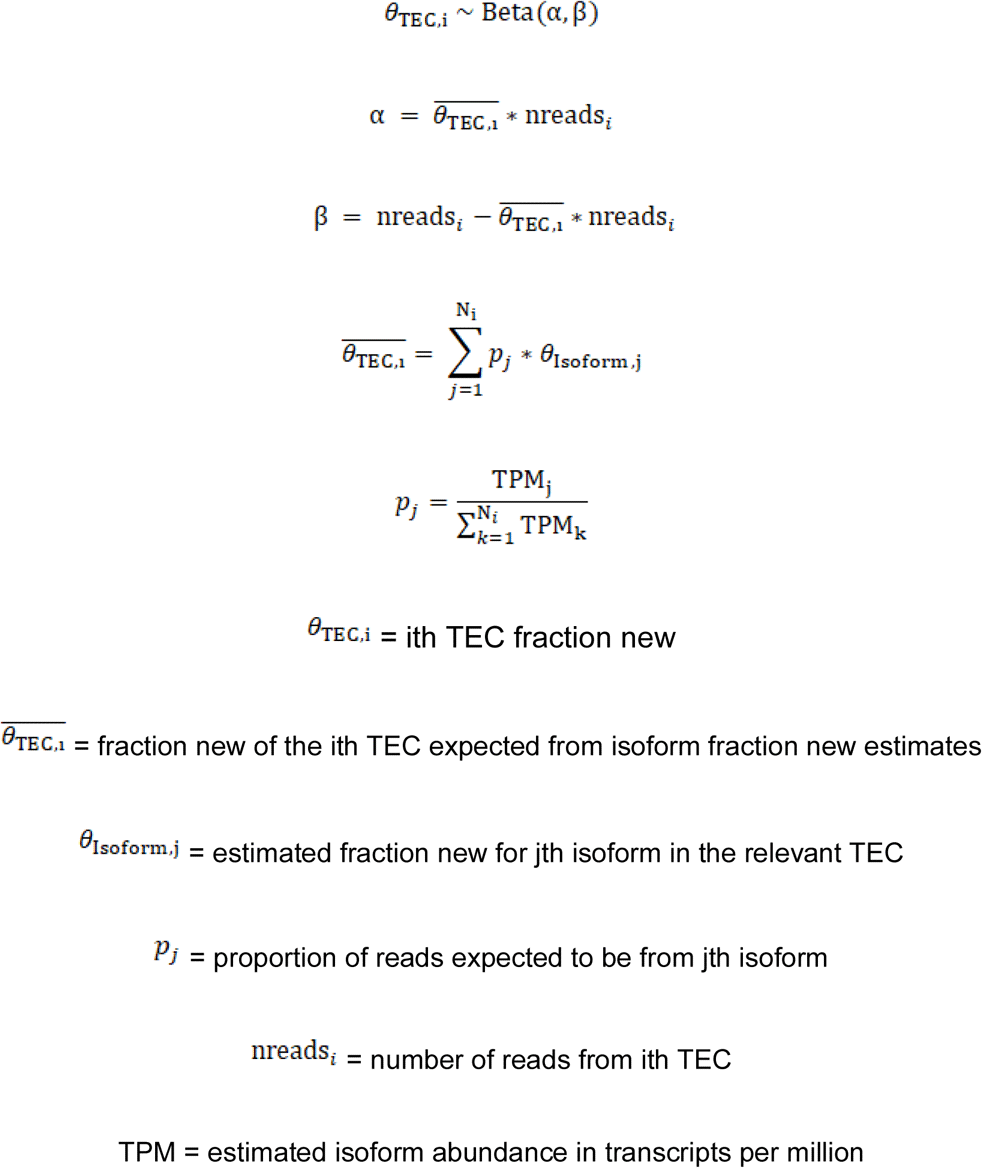

Uncertainty in isoform fraction new estimates are obtained from the Hessian matrix. Isoform degradation and synthesis rate constants are then estimated from isoform fraction news as follows:

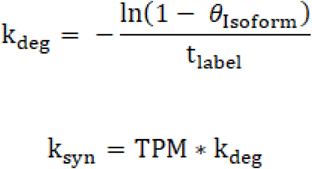

All isoform-level analyses in Figure 2 made use of the RefSeq annotation. Isoform analysis from Figure 4 onward used the Mix annotation from AnnotationCleaner.

### Simulating NR-seq data

NR-seq data was simulated using NRsim, a Snakemake pipeline we developed (https://github.com/isaacvock/NRsim). NRsim combines polyester’s simulation of standard RNA-seq data with custom scripting to add T-to-C mutations to simulated sequencing reads in accordance with simulated kinetic parameter distributions and label times [88]. The input to NRsim is a real RNA-seq FASTQ file (or pair of FASTQ files), an annotation of transcript isoforms, and a genome FASTA file. The real data is used to estimate abundances of isoforms in the provided annotation using Salmon [89]. A filtered annotation is generated, removing isoforms with an estimated TPM < 1, and polyester [88] is provided with this filtered annotation as well as the estimated isoform abundances. Kinetic parameters of synthesis and degradation are randomly drawn for each major transcript isoform (i.e., the most highly expressed isoform from each gene), with (by default) a 50% chance of differential isoform turnover explaining differences in alternative isoform abundance for the minor isoforms. Each read from a given isoform is then randomly assigned the status of “new” or “old” with probability equal to the simulated fraction new (calculated from the simulated degradation rate constant). Each T in a new read has a user - specified probability of being converted to a C, referred to as the “pnew”. The final FASTQ files are shuffled so as to not interfere with isoform abundance parameter estimation via tools like RSEM.

For the simulations in **Figures 1** and **S1**, the RefSeq annotation and a SMG1i treated replicate were provided to NRsim. For the simulations in **Figure S4**, RefSeq trimmed by AnnotationCleaner was provided instead, so that we could test analyzing the simulated data with the untrimmed and unfiltered RefSeq annotation.

### Isoform Decay Range (IDR) analysis

The isoform degradation range (IDR) was determined using the output of EZbakR’s AverageAndRegularize() function. For each multi-isoform gene, IDR was calculated as the difference in the replicate average log(k_deg_) between the least and most stable isoform under a given condition (DMSO or SMG1i) (**Table S4**). Statistical significance was assessed using a two-tailed z-test, comparing the ratio of the range to the total uncertainty in the range against a standard normal distribution. To compare IDRs between DMSO- and SMG1i-treated cells, the IDR difference served as the test statistic numerator, while the denominator represented the total uncertainty of this difference. p-values were adjusted for multiple testing using the Benjamini-Hochberg method.

### Conserved poison exon analysis

Potential alternative splicing events were extracted from the Mix annotation GTF using the SUPPA2 generateEvents function [90]. Junctions evaluated by EZbakR were assigned to SUPPA splicing events and integrated with factR2 PTC predictions [83]. Coordinates corresponding to inclusion and exclusion junctions of conserved poison exons were obtained from [50] and converted to hg38 coordinates using the UCSC liftover tool [91]. Conserved poison exons selected for further analysis were those that had exact matches to at least one inclusion and one exclusion coordinate of SUPPA2-identified splicing events and were predicted to contain PTCs by factR2 [83].

### PARE-seq reanalysis

Reads from the GSE61398 [56] dataset were trimmed using fastp [92] [-L -b 20]. The trimmed reads were then aligned to the hg38 reference genome using STAR [93] [--sjdbOverhang 19]. PARE peaks were called and read coverage was quantified using the CageFightR [94] package, with a coverage cutoff of 1 TPM per cluster and a cluster merge distance of 20 nucleotides. The Degust [95] server was used to TMM normalize cluster coverage. Differential coverage was evaluated using RUV-seq [96], applying the RUVr option (k=1). The highest coverage positions within each cluster were mapped to the Mix transcript annotation coordinates using the GenomicFeatures [97] and RCAS [98] packages. Clusters mapping to positions between −10 and +50 relative to TCs and significantly depleted in SMG6 or UPF1 knockdown conditions (L2FC < −1 and FDR < 0.05) were selected for further visual inspection.

### XRN1 and SMG6 knockdowns and Akron5-Seq library preparation

For depletion of XRN1 and SMG6, HEK-293T cells were seeded at 3 × 10⁵ cells per well and reverse transfected with 40 nM siRNA targeting *XRN1* (Forward: 5’ AGAUGAACUUACCGUAGAA 3’; Reverse: 5’ UUCUACGGUAAGUUCAUCU 3’) or *SMG6* (Forward: 5’ GCUGCAGGUUACUUACAAG 3’; Reverse: 5’ CUUGUAAGUAACCUGCAGC 3’) [99], or a *negative control* (Negative Control #2, Thermo Scientific) using Lipofectamine RNAiMAX per manufacturer instructions. After 48 hours, cells were replated at 5 × 10⁵ cells per well in a 6 - well plate. The next day, total RNA was extracted using TRIzol™ (Invitrogen) with overnight isopropanol precipitation at −80°C. RNA was then DNase-treated (dsDNase, Thermo Scientific) and purified with 0.6× SPRIselect beads (Beckman Coulter).

Three biological replicates were processed per condition. A total of 150 µg RNA per sample was used to generate Akron libraries, as described previously [55], with modifications. Total RNA underwent two rounds of poly(A) selection using the NEB Magnetic mRNA Isolation Kit and 100 µL oligo(dT) beads. RNA was denatured for 90 sec at 80°C, incubated with beads for 15 min at room temperature, and sequentially washed with 1× Bind/Wash Buffer (5 mM Tris-HCl pH 7.5, 0.5 mM EDTA, 1 M NaCl), followed by Wash Buffer B (10 mM Tris-HCl pH 7.5, 1 mM EDTA, 150 mM LiCl) and nuclease-free water.

Poly(A)-enriched mRNA was ligated to a 0.2 µM 5’ biotin-tagged RNA adaptor using T4 RNA Ligase (Thermo Scientific) at 16°C, 800 rpm, overnight. The following day, after three washes with Bind/Wash buffer, mRNA was fragmented using 10× Fragmentation Reagent (Thermo Scientific) for 7 min at 70 °C. Fragmented mRNA was purified using Dynabeads M-280 streptavidin beads (Invitrogen) in a 1:1 ratio for 30 min at room temperature. After washing, end repair was performed for 1 h at 37 °C using T4 PNK (NEB). The mRNA was then ligated to a 0.1 µM 3’ adaptor at 16°C, 800 rpm, overnight. The 3’ adaptor contained a 5ʹ phosphate group to enhance ligation efficiency, a 3ʹ-dideoxycytidine to prevent unwanted ligation, and a 10-nucleotide unique molecular identifier (UMI) for PCR deduplication.

cDNA synthesis was performed using SuperScript III Reverse Transcriptase (Invitrogen) and an RT primer (sequence listed in table below), followed by two rounds of PCR amplification using NEBNext UltraII Q5 MM. The first PCR used DP5 and RT primers (16 cycles), while the second PCR used the 5’ SR universal primer and NEB index primers (8 cycles) under the following conditions: 98°C for 30 s, followed by cycles of 98°C for 10 sec and 65°C for 75 sec, and a final extension at 65°C for 5 min. PCR products were ethanol-precipitated overnight at −80°C with 2.5X ethanol, 0.1X 3M sodium acetate pH 5.5, and GlycoBlue. Size selection was performed using Pippin Prep (1.5% agarose, marker K, Sage Science) to isolate fragments between 500–800 bp. Final libraries were quantified using the Quant-IT PicoGreen dsDNA Kit (Promega), analyzed on an Agilent Tapestation 2200 with DS1000 screentape, and sequenced on an Illumina NovaSeq X plus at a depth of >30 million reads per library.

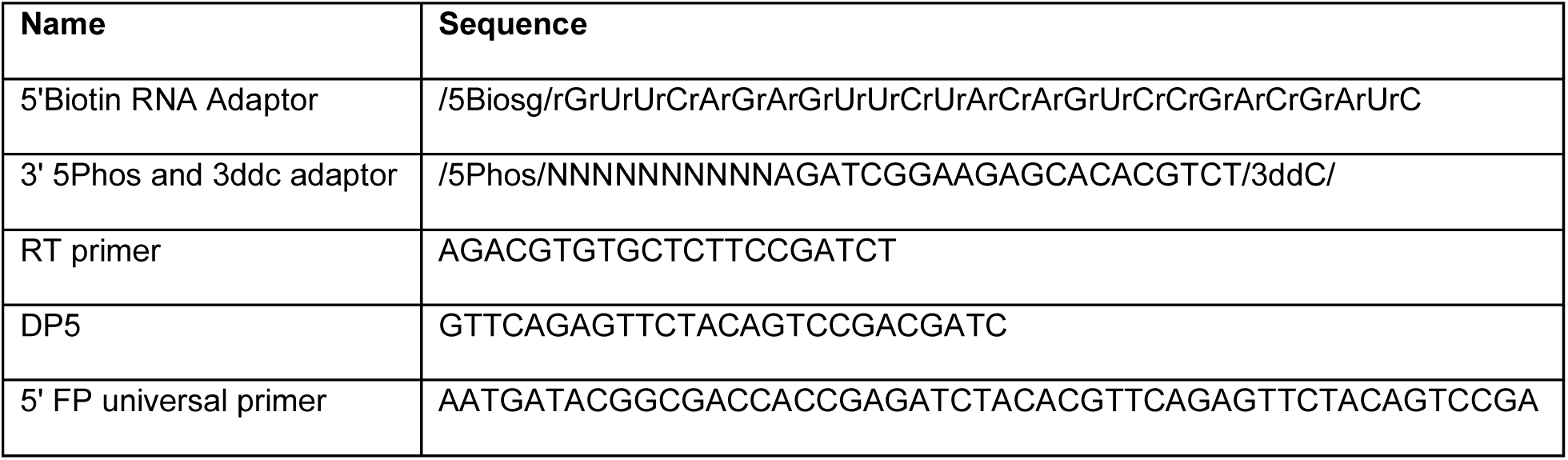

### Akron5-Seq data processing and analysis

UMIs were extracted from sequencing reads using UMI-tools [100]. Reads were subsequently quality-trimmed using fastp [92] with automatic adapter detection and poly-G tail removal. Trimmed reads were mapped to Repbase [101] using STAR [93] (v2.7.6a) with up to 30 allowed multimapping locations. Unmapped reads were subsequently aligned to the hg38 reference genome using STAR [93] with parameters optimized for accurate splice junction detection and paired-end overlap resolution. Uniquely mapping reads were extracted using samtools [80] (q=255) and deduplicated based on unique molecular identifiers (UMIs) with UMI-tools [100]. Only properly paired, non-chimeric alignments were retained. To quantify putative SMG6 cleavage sites, 5’ end read alignments were extracted from deduplicated BAM files using samtools [80] and converted into BigWig tracks with CAGEfightR [94]. Cleavage clusters were identified using CAGEfightR [94] with a threshold of 1 TPM and merging distances of 20 nucleotides. Cleavage site counts were assigned to transcript features using the Mix annotation and exported. Differential cluster abundance analysis was performed using Degust [95]. Cleavage clusters with >10 counts in at least three samples were normalized with TMM, and batch effects were corrected using RUVr [96] (k=2). Differentially abundant clusters were identified with edgeR maximum likelihood estimation. Clusters mapping to positions between −10 and +50 relative to non-PTC isoform TCs, significantly depleted in SMG6 knockdown conditions (L2FC < −1 and FDR < 0.05), and significantly stabilized by SMG1i (L2FC(k_deg_) < −0.25, padj < 0.05) were selected for further visual inspection. See **TableS5** for analysis output.

### LASSO regression and classification models

The glmnet R package was used to train LASSO regression models with our data [102,103]. We trained two types of models in this work: a degradation rate constant regression model and a SMG1i sensitivity classification model. In both cases, feature engineering was performed to make LASSO regression coefficients comparable across features. Numeric features were standardized (subtracted the mean and divided by the standard deviation), with numeric features ranging many orders of base-10 magnitude being log_10_ transformed prior to standardization. Engineering categorical features is more difficult. We settled on a strategy that involved ordering categorical features based on their average outcome variable value (e.g., lowest to highest degradation rate constant). For categorical features with more than 10 groups, we pooled groups with similar average outcome variables. This is to avoid artificially strong correlation between the ranks of a categorical feature and the outcome due to super-fine stratification of the data. We then z-scored the ranks assigned to each group and used that as the engineered feature. While this approach still risks artificially inflating the importance assigned to categorical features, we note that categorical features were never identified as the strongest predictors in any of our models. We also only used feature importances as inspiration for downstream analyses to more carefully probe the impact that these features may be having.

For the degradation rate constant regression model, we only considered transcript isoforms with high confidence estimates. Thus, we filtered out low coverage isoforms, isoforms with high degradation rate constant estimate uncertainty, and isoforms flagged by either of our filtering criterion (**Figure 3A**). We then trained the model with 100 different bootstrapped samples of the engineered features and DMSO treated degradation rate constant estimates, to assess stability of estimated feature importances. An optimal lambda value was determined through 10-fold cross validation on each bootstrapped dataset.

For the SMG1i sensitivity classification model, we only considered transcript isoforms which we could confidently assign the status of SMG1i sensitive or not. Thus, we filtered out low coverage isoforms and isoforms with high degradation rate constant estimate uncertainty. High confidence SMG1i-sensitive isoforms were the remaining isoforms that had a log_2_-fold change in their degradation rate constant of less than −0.7, and with a degradation rate constant after SMG1i inhibition less than 65% of its value prior to SMG1i inhibition (suggesting at least around 35% of its degradation is attributable to NMD). High confidence SMG1i insensitive isoforms were those with a log_2_-fold change in their degradation rate constant of around 0 (or that had estimated increases in their rate constant). For the classification model, we did not filter out unsupported isoforms or those from high JCC genes, as we were using a reduced set of isoforms to begin with and suspected that comparative analyses of degradation rate constant estimates were more robust to annotation discrepancies than single-condition estimates. To deal with the large class imbalance, we downsampled the insensitive isoform set to the size of the sensitive set. Feature importances were estimated for 100 different downsamplings to assess stability of these estimates. An optimal lambda value was determined through 10-fold cross validation on each downsampled dataset.

Features used in these analyses are provided in **TableS6**. Briefly, sequence features were calculated from the Mix annotation file using custom R and Python scripts. Additional data, such as RF1 and RF2 peptide release rates [104] and codon optimality [105], were either directly merged or processed before integration, as detailed below.

### CLIP-UPF1 affinity purification-seq reanalysis

Reads from the GSE134059 [61] dataset were trimmed using fastp [92] with parameters [-L -b 20]. The trimmed reads were then aligned to the hg38 reference genome and Mix transcript annotation using default STAR [93] parameters. Isoform quantification was performed using RSEM [19], and differential expression analysis of CLIP-UPF1 vs GFP AP was conducted via the Degust [95] server.

### TC-containing exon PhastCons scoring

To identify exons containing stop codons, a GTF file was processed using the R package GenomicRanges [97]. Exon entries were filtered, and stop codon positions were determined from the Mix transcript annotation. Exons overlapping stop codons were extracted and converted to BED format. PhastCons conservation scores were then obtained from the UCSC hg38 100-way alignment using the phastCons100way.UCSC.hg38 (Bioconductor) package. The BED file was imported as a GenomicRanges object, and PhastCons scores were computed for the selected exons.

### Isoform Kozak sequence scoring

A custom python script was used to assess the strength of Kozak sequences in the Mix transcript annotation FASTA file. A position-specific scoring matrix [106,107] was defined based on nucleotide positions within the 10-base Kozak sequence. For each sequence in the FASTA file, a Kozak score was computed by summing the matrix values for the corresponding nucleotides at each position.

## Supplemental information

Document S1. Figures S1–S7

Table S1. Excel file containing STAR alignment and EZbakR labeling information.

Table S2. Excel file containing EZbakR Isoform-level outputs.

Table S3. Excel file containing EZbakR Junction-level output.

Table S4. Excel file containing isoform decay range (IDR) estimates.

Table S5. Excel file containing analyzed Akron-seq output.

Table S6. Excel file containing features considered in the LASSO regression models.

## Declarations

### Lead contact

Requests for further information and resources should be directed to and will be fulfilled by the lead contact, J. Robert Hogg.

### Availability of data and materials

Short-read NR-seq FASTQ and processed bedgraph files and long-read BAM files have been deposited at GEO as GSE289489, and short-read Akron5-seq FASTQ and processed BigWig files have been deposited at GEO as GSE291379. The data are publicly available as of the date of publication. All original code has been deposited on GitHub and is publicly available at the date of publication. Any additional information required to reanalyze the data reported in this paper is available from the lead contact upon request. [https://github.com/isaacvock/EZbakR_Isoforms_and_NMD_scripts]

### Author contributions

Conceptualization I.W.V., J.W.M., M.M., J.R.H., M.D.S

Data curation I.W.V., J.W.M.

Formal analysis I.W.V., J.W.M.

Funding acquisition J.I., J.R.H., M.D.S.

Investigation J.W.M., I.W.V., N.H., P.T., M.M., A.Z.

Resources G.R., I.N-M.L., J.I.

Methodology I.W.V. Software I.W.V.

Supervision J.R.H., M.D.S., J.I.

Visualization J.W.M., I.W.V., J.R.H.

Writing-original draft J.W.M., I.W.V., J.R.H.

Writing-reviewing and editing J.W.M., I.W.V., M.M., G.R., I.N-M.L., J.I., N.H., P.T., A.Z., J.R.H., M.D.S.

## Acknowledgments

We thank members of the Hogg and Simon Labs for helpful discussions. This work utilized the resources of the NIH HPC Biowulf cluster (https://hpc.nih.gov).

## References

1. Karousis ED, Mühlemann O. Nonsense-Mediated mRNA Decay Begins Where Translation Ends. Cold Spring Harb Perspect Biol. 2019;11:a032862.

2. Kurosaki T, Popp MW, Maquat LE. Quality and quantity control of gene expression by nonsense-mediated mRNA decay. Nat Rev Mol Cell Biol. 2019;20:406–20.

3. McMahon M, Maquat LE. Exploring the therapeutic potential of modulating nonsense-mediated mRNA decay. RNA. 2025;31:333–48.

4. Rebbapragada I, Lykke-Andersen J. Execution of nonsense-mediated mRNA decay: what defines a substrate? Curr Opin Cell Biol. 2009;21:394–402.

5. Nagy E, Maquat LE. A rule for termination-codon position within intron-containing genes: when nonsense affects RNA abundance. Trends Biochem Sci. 1998;23:198–9.

6. Le Hir H, Gatfield D, Izaurralde E, Moore MJ. The exon-exon junction complex provides a binding platform for factors involved in mRNA export and nonsense-mediated mRNA decay. EMBO J. 2001;20:4987–97.

7. Kim VN, Kataoka N, Dreyfuss G. Role of the nonsense-mediated decay factor hUpf3 in the splicing-dependent exon-exon junction complex. Science. 2001;293:1832–6.

8. Lindeboom RGH, Supek F, Lehner B. The rules and impact of nonsense-mediated mRNA decay in human cancers. Nat Genet. 2016;48:1112–8.

9. Kim Y-G, Kang H, Lee B, Jang H-J, Park J-H, Ha C, et al. A spectrum of nonsense-mediated mRNA decay efficiency along the degree of mutational constraint. Commun Biol. 2024;7:1461.

10. Lindeboom RGH, Vermeulen M, Lehner B, Supek F. The impact of nonsense-mediated mRNA decay on genetic disease, gene editing and cancer immunotherapy. Nat Genet. 2019;51:1645–51.

11. Supek F, Lehner B, Lindeboom RGH. To NMD or not to NMD: Nonsense-mediated mRNA decay in cancer and other genetic diseases. Trends Genet. 2021;37:657–68.

12. Muñoz O, Lore M, Jagannathan S. The long and short of EJC-independent nonsense-mediated RNA decay. Biochem Soc Trans. 2023;51:1121–9.

13. Schofield JA, Duffy EE, Kiefer L, Sullivan MC, Simon MD. TimeLapse-seq: adding a temporal dimension to RNA sequencing through nucleoside recoding. Nat Methods. 2018;15:221–5.

14. Herzog VA, Reichholf B, Neumann T, Rescheneder P, Bhat P, Burkard TR, et al. Thiol-linked alkylation of RNA to assess expression dynamics. Nat Methods. 2017;14:1198–204.

15. Riml C, Amort T, Rieder D, Gasser C, Lusser A, Micura R. Osmium-mediated transformation of 4-thiouridine to cytidine as key to study RNA dynamics by sequencing. Angew Chem Int Ed Engl. 2017;56:13479–83.

16. Duffy EE, Schofield JA, Simon MD. Gaining insight into transcriptome-wide RNA population dynamics through the chemistry of 4-thiouridine. Wiley Interdiscip Rev RNA. 2019;10:e1513.

17. Vock IW, Mabin JW, Machyna M, Zhang A, Hogg JR, Simon MD. Expanding and improving analyses of nucleotide recoding RNA-seq experiments with the EZbakR suite. bioRxiv. 2024. p. 2024.10.14.617411.

18. Jürges C, Dölken L, Erhard F. Dissecting newly transcribed and old RNA using GRAND-SLAM. Bioinformatics. 2018;34:i218–26.

19. Li B, Dewey CN. RSEM: accurate transcript quantification from RNA-Seq data with or without a reference genome. BMC Bioinformatics. 2011;12:323.

20. Kishor A, Fritz SE, Hogg JR. Nonsense-mediated mRNA decay: The challenge of telling right from wrong in a complex transcriptome. Wiley Interdiscip Rev RNA. 2019;10:e1548.

21. Gopalsamy A, Bennett EM, Shi M, Zhang W-G, Bard J, Yu K. Identification of pyrimidine derivatives as hSMG-1 inhibitors. Bioorg Med Chem Lett. 2012;22:6636–41.

22. Ni JZ, Grate L, Donohue JP, Preston C, Nobida N, O’Brien G, et al. Ultraconserved elements are associated with homeostatic control of splicing regulators by alternative splicing and nonsense-mediated decay. Genes Dev. 2007;21:708–18.

23. Lareau LF, Inada M, Green RE, Wengrod JC, Brenner SE. Unproductive splicing of SR genes associated with highly conserved and ultraconserved DNA elements. Nature. 2007;446:926–9.

24. Narain A, Bhandare P, Adhikari B, Backes S, Eilers M, Dölken L, et al. Targeted protein degradation reveals a direct role of SPT6 in RNAPII elongation and termination. Mol Cell. 2021;81:3110–27.e14.

25. Xiao R, Tang P, Yang B, Huang J, Zhou Y, Shao C, et al. Nuclear matrix factor hnRNP U/SAF-A exerts a global control of alternative splicing by regulating U2 snRNP maturation. Mol Cell. 2012;45:656–68.

26. Haimovich G, Medina DA, Causse SZ, Garber M, Millán-Zambrano G, Barkai O, et al. Gene expression is circular: factors for mRNA degradation also foster mRNA synthesis. Cell. 2013;153:1000–11.

27. Berry S, Müller M, Rai A, Pelkmans L. Feedback from nuclear RNA on transcription promotes robust RNA concentration homeostasis in human cells. Cell Syst. 2022;13:454– 70.e15.

28. Britto-Borges T, Gehring NH, Boehm V, Dieterich C. NMDtxDB: data-driven identification and annotation of human NMD target transcripts. RNA. 2024;30:1277–91.

29. Karousis ED, Gypas F, Zavolan M, Mühlemann O. Nanopore sequencing reveals endogenous NMD-targeted isoforms in human cells. Genome Biol. 2021;22:223.

30. Chousal JN, Sohni A, Vitting-Seerup K, Cho K, Kim M, Tan K, et al. Progression of the pluripotent epiblast depends upon the NMD factor UPF2. Development. 2022;149:dev200764.

31. Bicknell AA, Cenik C, Chua HN, Roth FP, Moore MJ. Introns in UTRs: why we should stop ignoring them. Bioessays. 2012;34:1025–34.

32. Chabbert CD, Eberhart T, Guccini I, Krek W, Kovacs WJ. Correction of gene model annotations improves isoform abundance estimates: the example of ketohexokinase (*Khk*). F1000Res. 2019;7:1956.

33. Varabyou A, Salzberg SL, Pertea M. Effects of transcriptional noise on estimates of gene and transcript expression in RNA sequencing experiments. Genome Res. 2021;31:301–8.

34. Pertea M, Pertea GM, Antonescu CM, Chang T-C, Mendell JT, Salzberg SL. StringTie enables improved reconstruction of a transcriptome from RNA-seq reads. Nat Biotechnol. 2015;33:290–5.

35. Kovaka S, Zimin AV, Pertea GM, Razaghi R, Salzberg SL, Pertea M. Transcriptome assembly from long-read RNA-seq alignments with StringTie2. Genome Biol. 2019;20:278.

36. Soneson C, Love MI, Patro R, Hussain S, Malhotra D, Robinson MD. A junction coverage compatibility score to quantify the reliability of transcript abundance estimates and annotation catalogs. Life Sci Alliance. 2019;2:e201800175.

37. Love MI, Hogenesch JB, Irizarry RA. Modeling of RNA-seq fragment sequence bias reduces systematic errors in transcript abundance estimation. Nat Biotechnol. 2016;34:1287–91.

38. O’Leary NA, Wright MW, Brister JR, Ciufo S, Haddad D, McVeigh R, et al. Reference sequence (RefSeq) database at NCBI: current status, taxonomic expansion, and functional annotation. Nucleic Acids Res. 2016;44:D733–45.

39. Harrison PW, Amode MR, Austine-Orimoloye O, Azov AG, Barba M, Barnes I, et al. Ensembl 2024. Nucleic Acids Res. 2024;52:D891–9.

40. He PC, Wei J, Dou X, Harada BT, Zhang Z, Ge R, et al. Exon architecture controls mRNA m6A suppression and gene expression. Science. 2023;379:677–82.

41. Yang X, Triboulet R, Liu Q, Sendinc E, Gregory RI. Exon junction complex shapes the m6A epitranscriptome. Nat Commun. 2022;13:7904.

42. Uzonyi A, Dierks D, Nir R, Kwon OS, Toth U, Barbosa I, et al. Exclusion of m6A from splice-site proximal regions by the exon junction complex dictates m6A topologies and mRNA stability. Mol Cell. 2023;83:237–51.e7.

43. Agarwal V, Kelley DR. The genetic and biochemical determinants of mRNA degradation rates in mammals. Genome Biol. 2022;23:245.

44. Lugowski A, Nicholson B, Rissland OS. DRUID: a pipeline for transcriptome-wide measurements of mRNA stability. RNA. 2018;24:623–32.

45. Pereira FJC, Teixeira A, Kong J, Barbosa C, Silva AL, Marques-Ramos A, et al. Resistance of mRNAs with AUG-proximal nonsense mutations to nonsense-mediated decay reflects variables of mRNA structure and translational activity. Nucleic Acids Res. 2015;43:6528–44.

46. Neu-Yilik G, Amthor B, Gehring NH, Bahri S, Paidassi H, Hentze MW, et al. Mechanism of escape from nonsense-mediated mRNA decay of human beta-globin transcripts with nonsense mutations in the first exon. RNA. 2011;17:843–54.

47. Russell PJ, Slivka JA, Boyle EP, Burghes AHM, Kearse MG. Translation reinitiation after uORFs does not fully protect mRNAs from nonsense-mediated decay. RNA. 2023;29:735–44.

48. Hoek TA, Khuperkar D, Lindeboom RGH, Sonneveld S, Verhagen BMP, Boersma S, et al. Single-Molecule Imaging Uncovers Rules Governing Nonsense-Mediated mRNA Decay. Mol Cell. 2019;75:324–39.e11.

49. Karczewski KJ, Francioli LC, Tiao G, Cummings BB, Alföldi J, Wang Q, et al. The mutational constraint spectrum quantified from variation in 141,456 humans. Nature. 2020;581:434–43.

50. Thomas JD, Polaski JT, Feng Q, De Neef EJ, Hoppe ER, McSharry MV, et al. RNA isoform screens uncover the essentiality and tumor-suppressor activity of ultraconserved poison exons. Nat Genet. 2020;52:84–94.

51. Yepiskoposyan H, Aeschimann F, Nilsson D, Okoniewski M, Mühlemann O. Autoregulation of the nonsense-mediated mRNA decay pathway in human cells. RNA. 2011;17:2108–18.

52. Huang L, Lou C-H, Chan W, Shum EY, Shao A, Stone E, et al. RNA homeostasis governed by cell type-specific and branched feedback loops acting on NMD. Mol Cell. 2011;43:950–61.

53. Singh G, Rebbapragada I, Lykke-Andersen J. A competition between stimulators and antagonists of Upf complex recruitment governs human nonsense-mediated mRNA decay. PLoS Biol. 2008;6:e111.

54. Karam R, Lou C-H, Kroeger H, Huang L, Lin JH, Wilkinson MF. The unfolded protein response is shaped by the NMD pathway. EMBO Rep. 2015;16:599–609.

55. Ibrahim F, Mourelatos Z. Capturing 5’ and 3’ native ends of mRNAs concurrently with Akron sequencing. Nat Protoc. 2019;14:1578–602.

56. Schmidt SA, Foley PL, Jeong D-H, Rymarquis LA, Doyle F, Tenenbaum SA, et al. Identification of SMG6 cleavage sites and a preferred RNA cleavage motif by global analysis of endogenous NMD targets in human cells. Nucleic Acids Res. 2015;43:309–23.

57. Lykke-Andersen S, Chen Y, Ardal BR, Lilje B, Waage J, Sandelin A, et al. Human nonsense-mediated RNA decay initiates widely by endonucleolysis and targets snoRNA host genes. Genes Dev. 2014;28:2498–517.

58. Ottens F, Boehm V, Sibley CR, Ule J, Gehring NH. Transcript-specific characteristics determine the contribution of endo- and exonucleolytic decay pathways during the degradation of nonsense-mediated decay substrates. RNA. 2017;23:1224–36.

59. Fair B, Buen Abad Najar CF, Zhao J, Lozano S, Reilly A, Mossian G, et al. Global impact of unproductive splicing on human gene expression. Nat Genet. 2024;56:1851–61.

60. Fritz SE, Ranganathan S, Wang CD, Hogg JR. An alternative UPF1 isoform drives conditional remodeling of nonsense-mediated mRNA decay. EMBO J. 2022;41:e108898.

61. Kishor A, Fritz SE, Haque N, Ge Z, Tunc I, Yang W, et al. Activation and inhibition of nonsense-mediated mRNA decay control the abundance of alternative polyadenylation products. Nucleic Acids Res. 2020;48:7468–82.

62. Hogg JR, Goff SP. Upf1 senses 3’UTR length to potentiate mRNA decay. Cell. 2010;143:379–89.

63. Johns L, Grimson A, Kuchma SL, Newman CL, Anderson P. Caenorhabditis elegans SMG-2 selectively marks mRNAs containing premature translation termination codons. Mol Cell Biol. 2007;27:5630–8.

64. Colombo M, Karousis ED, Bourquin J, Bruggmann R, Mühlemann O. Transcriptome-wide identification of NMD-targeted human mRNAs reveals extensive redundancy between SMG6- and SMG7-mediated degradation pathways. RNA. 2017;23:189–201.

65. Imamachi N, Salam KA, Suzuki Y, Akimitsu N. A GC-rich sequence feature in the 3’ UTR directs UPF1-dependent mRNA decay in mammalian cells. Genome Res. 2017;27:407–18.

66. Kolakada D, Campbell AE, Galvis LB, Li Z, Lore M, Jagannathan S. A system of reporters for comparative investigation of EJC-independent and EJC-enhanced nonsense-mediated mRNA decay. Nucleic Acids Res. 2024;52:e34.

67. Li G, Rice CM. The signal for translational readthrough of a UGA codon in Sindbis virus RNA involves a single cytidine residue immediately downstream of the termination codon. J Virol. 1993;67:5062–7.

68. McCaughan KK, Brown CM, Dalphin ME, Berry MJ, Tate WP. Translational termination efficiency in mammals is influenced by the base following the stop codon. Proc Natl Acad Sci U S A. 1995;92:5431–5.

69. Skuzeski JM, Nichols LM, Gesteland RF, Atkins JF. The signal for a leaky UAG stop codon in several plant viruses includes the two downstream codons. J Mol Biol. 1991;218:365–73.

70. Bonetti B, Fu L, Moon J, Bedwell DM. The efficiency of translation termination is determined by a synergistic interplay between upstream and downstream sequences in Saccharomyces cerevisiae. J Mol Biol. 1995;251:334–45.

71. Daly CWP, Kristjánsdóttir K, West JD, Vu LT, Fogarty EA, Geissler R, et al. A novel regulatory pathway recognizes and degrades transcripts with long 3′ untranslated regions. bioRxiv. 2024. p. 2024.03.11.584429.

72. Kishor A, Ge Z, Hogg JR. hnRNP L-dependent protection of normal mRNAs from NMD subverts quality control in B cell lymphoma. EMBO J. 2019;38:e99128.

73. Ge Z, Quek BL, Beemon KL, Hogg JR. Polypyrimidine tract binding protein 1 protects mRNAs from recognition by the nonsense-mediated mRNA decay pathway. Elife [Internet]. 2016 [cited 2025 Feb 10];5. Available from: https://pubmed.ncbi.nlm.nih.gov/26744779/

74. Luo Y, Schofield JA, Simon MD, Slavoff SA. Global profiling of cellular substrates of human Dcp2. Biochemistry. 2020;59:4176–88.

75. Ietswaart R, Smalec BM, Xu A, Choquet K, McShane E, Jowhar ZM, et al. Genome-wide quantification of RNA flow across subcellular compartments reveals determinants of the mammalian transcript life cycle. Mol Cell. 2024;84:2765–84.e16.

76. Simon M, Schofield J, Duffy E, Kiefer L, Sullivan M, Simon M. Protocol for TimeLapse-seq. Protoc Exch [Internet]. 2018 [cited 2025 Feb 5]; Available from: https://www.researchsquare.com/article/nprot-6521/latest.pdf

77. Baldwin A, Morris AR, Mukherjee N. An Easy, Cost-Effective, and Scalable Method to Deplete Human Ribosomal RNA for RNA-seq. Curr Protoc. 2021;1:e176.

78. Quinlan AR, Hall IM. BEDTools: a flexible suite of utilities for comparing genomic features. Bioinformatics. 2010;26:841–2.

79. Li H. Minimap2: pairwise alignment for nucleotide sequences. Bioinformatics. 2018;34:3094–100.

80. Li H, Handsaker B, Wysoker A, Fennell T, Ruan J, Homer N, et al. The Sequence Alignment/Map format and SAMtools. Bioinformatics. 2009;25:2078–9.

81. Liao Y, Smyth GK, Shi W. featureCounts: an efficient general purpose program for assigning sequence reads to genomic features. Bioinformatics. 2014;30:923–30.

82. Vock IW, Simon MD. bakR: uncovering differential RNA synthesis and degradation kinetics transcriptome-wide with Bayesian hierarchical modeling. RNA. 2023;29:958–76.

83. Zhuravskaya A, Yap K, Hamid F, Makeyev EV. Alternative splicing coupled to nonsense-mediated decay coordinates downregulation of non-neuronal genes in developing mouse neurons. Genome Biol. 2024;25:162.

84. Tardaguila M, de la Fuente L, Marti C, Pereira C, Pardo-Palacios FJ, Del Risco H, et al. SQANTI: extensive characterization of long-read transcript sequences for quality control in full-length transcriptome identification and quantification. Genome Res. 2018;28:396.

85. Tang S, Lomsadze A, Borodovsky M. Identification of protein coding regions in RNA transcripts. Nucleic Acids Res. 2015;43:e78.

86. Pertea G, Pertea M. GFF utilities: GffRead and GffCompare. F1000Res. 2020;9:304.

87. Varabyou A, Erdogdu B, Salzberg SL, Pertea M. Investigating open reading frames in known and novel transcripts using ORFanage. Nat Comput Sci. 2023;3:700–8.

88. Frazee AC, Jaffe AE, Langmead B, Leek JT. Polyester: simulating RNA-seq datasets with differential transcript expression. Bioinformatics. 2015;31:2778–84.

89. Patro R, Duggal G, Love MI, Irizarry RA, Kingsford C. Salmon provides fast and bias-aware quantification of transcript expression. Nat Methods. 2017;14:417–9.

90. Trincado JL, Entizne JC, Hysenaj G, Singh B, Skalic M, Elliott DJ, et al. SUPPA2: fast, accurate, and uncertainty-aware differential splicing analysis across multiple conditions. Genome Biology. 2018;19:1–11.

91. Hinrichs AS, Karolchik D, Baertsch R, Barber GP, Bejerano G, Clawson H, et al. The UCSC Genome Browser Database: update 2006. Nucleic Acids Res. 2006;34:D590–8.

92. Chen S, Zhou Y, Chen Y, Gu J. fastp: an ultra-fast all-in-one FASTQ preprocessor. Bioinformatics. 2018;34:i884–90.

93. Dobin A, Davis CA, Schlesinger F, Drenkow J, Zaleski C, Jha S, et al. STAR: ultrafast universal RNA-seq aligner. Bioinformatics. 2013;29:15–21.

94. Thodberg M, Thieffry A, Vitting-Seerup K, Andersson R, Sandelin A. CAGEfightR: analysis of 5’-end data using R/Bioconductor. BMC Bioinformatics. 2019;20:487.

95. Powell D. drpowell/degust 4.1.1 [Internet]. Zenodo; 2019. Available from: 10.5281/ZENODO.3258932

96. Risso D, Ngai J, Speed TP, Dudoit S. Normalization of RNA-seq data using factor analysis of control genes or samples. Nat Biotechnol. 2014;32:896–902.

97. Lawrence M, Huber W, Pagès H, Aboyoun P, Carlson M, Gentleman R, et al. Software for computing and annotating genomic ranges. PLoS Comput Biol. 2013;9:e1003118.

98. Uyar B, Yusuf D, Wurmus R, Rajewsky N, Ohler U, Akalin A. RCAS: an RNA centric annotation system for transcriptome-wide regions of interest. Nucleic Acids Res. 2017;45:e91.

99. Durand S, Franks TM, Lykke-Andersen J. Hyperphosphorylation amplifies UPF1 activity to resolve stalls in nonsense-mediated mRNA decay. Nat Commun. 2016;7:12434.

100. Smith T, Heger A, Sudbery I. UMI-tools: modeling sequencing errors in Unique Molecular Identifiers to improve quantification accuracy. Genome Res. 2017;27:491–9.

101. Bao W, Kojima KK, Kohany O. Repbase Update, a database of repetitive elements in eukaryotic genomes. Mob DNA. 2015;6:11.

102. Tay JK, Narasimhan B, Hastie T. Elastic net regularization paths for all generalized linear models. J Stat Softw. 2023;106:1–31.

103. Friedman J, Hastie T, Tibshirani R. Regularization paths for generalized linear models via coordinate descent. J Stat Softw. 2010;33:1–22.

104. Pierson WE, Hoffer ED, Keedy HE, Simms CL, Dunham CM, Zaher HS. Uniformity of peptide release is maintained by methylation of release factors. Cell Rep. 2016;17:11–8.

105. Forrest ME, Pinkard O, Martin S, Sweet TJ, Hanson G, Coller J. Codon and amino acid content are associated with mRNA stability in mammalian cells. PLoS One. 2020;15:e0228730.

106. Popa A, Lebrigand K, Barbry P, Waldmann R. Pateamine A-sensitive ribosome profiling reveals the scope of translation in mouse embryonic stem cells. BMC Genomics. 2016;17:52.

107. Kim K-B, Park K, Kong EB. A method for identifying splice sites and translation start sites in human genomic sequences. J Biochem Mol Biol. 2002;35:513–7.

