## Supplemental Figures for "Uncovering the isoform-resolution kinetic landscape of nonsense-mediated mRNA decay with EZbakR"

#### Figure S1

(A)

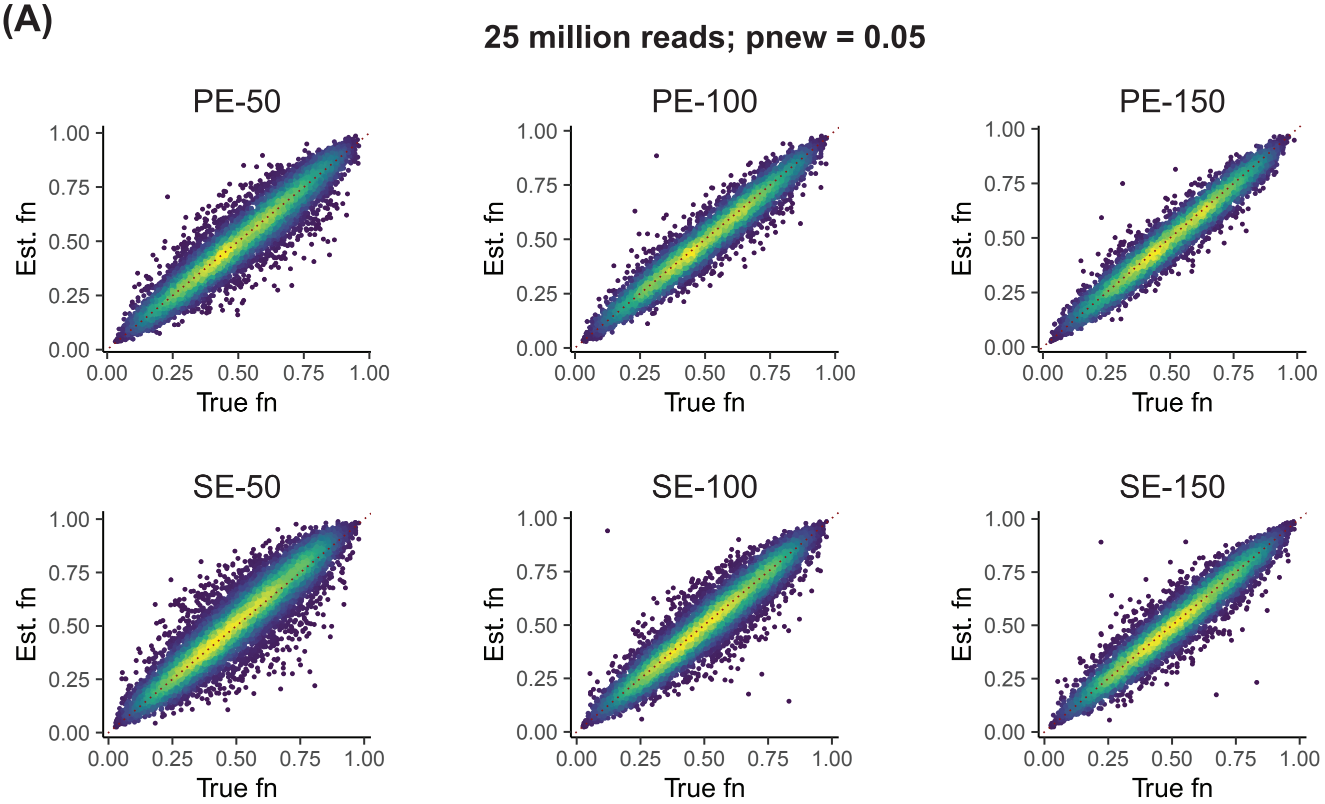
25 million reads; pnew = 0.05
PE-50
PE-100
PE-150
SE-50
SE-100
SE-150
Est. fn
True fn
0.00
0.25
0.50
0.75
1.00

(B)

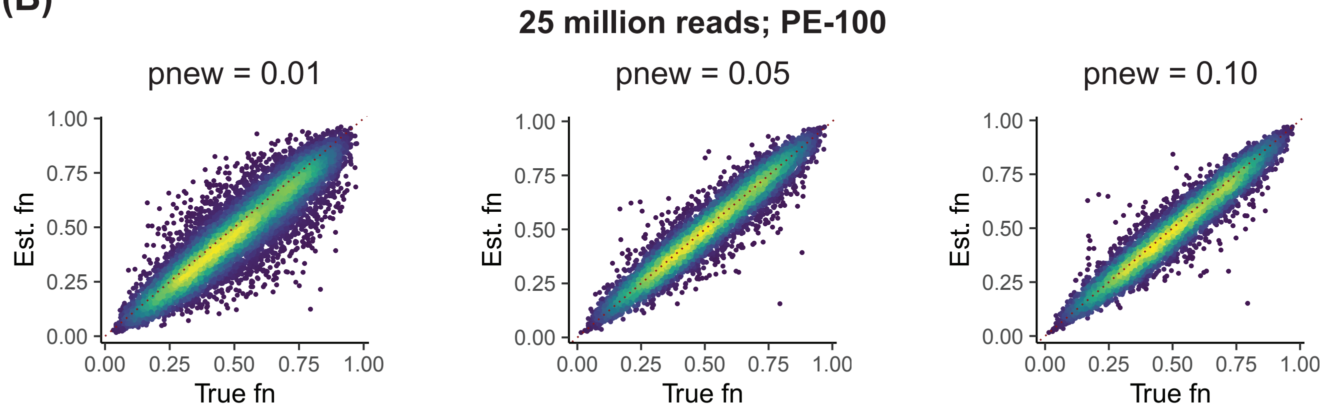
25 million reads; PE-100
pnew = 0.01
pnew = 0.05
pnew = 0.10
Est. fn
True fn
0.00
0.25
0.50
0.75
1.00

**Figure S1.** Associated with Figure 1

**(A)** Further simulated data assessment of isoform fraction new estimate accuracy across a range of read lengths. PE denotes paired-end data and SE denotes single-end data. Points are colored by density, and the red dotted line represents perfect estimation (y = x).

**(B)** Further simulated data assessment of isoform fraction new estimate accuracy across a range of incorporation rates (pnew; defined as the rate of T-to-C conversion in reads from new RNA). Points are colored by density, and the red dotted line represents perfect estimation (y = x).

#### Figure S2

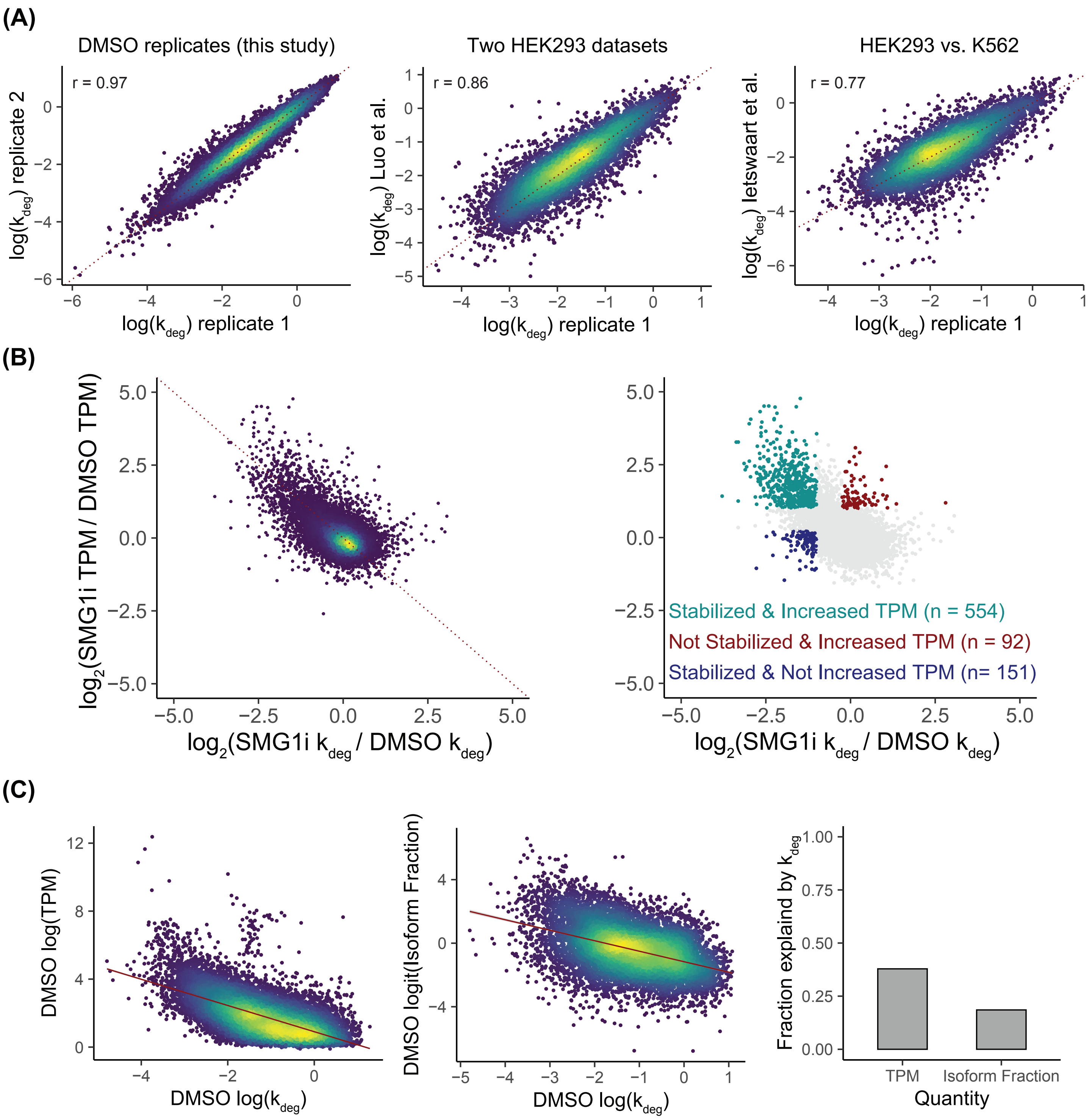
(A)
DMSO replicates (this study)
r = 0.97
log(k_deg) replicate 2
log(k_deg) replicate 1
Two HEK293 datasets
r = 0.86
log(k_deg) Luo et al.
log(k_deg) replicate 1
HEK293 vs. K562
r = 0.77
log(k_deg) letswaart et al.
log(k_deg) replicate 1
(B)
log2(SMG1i TPM / DMSO TPM)
log2(SMG1i k_deg / DMSO k_deg)
Stabilized & Increased TPM (n = 554)
Not Stabilized & Increased TPM (n = 92)
Stabilized & Not Increased TPM (n= 151)
log2(SMG1i k_deg / DMSO k_deg)
(C)
DMSO log(TPM)
DMSO log(k_deg)
DMSO logit(Isoform Fraction)
DMSO log(k_deg)
Fraction explaind by k_deg
TPM
Isoform Fraction
Quantity

**Figure S2.** Associated with Figure 2

**(A)** Consistency of EZbakR isoform-level degradation rate constant estimates. **Left:** comparison of two replicates of DMSO treated HEK293 cell degradation rate constant estimates from this study. **Middle:** comparison of one replicate of DMSO treated HEK293 cell degradation rate constant estimates from this study to those from one replicate of WT HEK293 cell from a previously published dataset [74]. **Right:** comparison of one replicate of DMSO treated HEK293 cell degradation rate constant estimates from this study to those from one replicate of WT K562 cells from a previously published dataset [75].

**(B)** Extent to which differential stability explains differential expression upon SMG1 inhibition. Both plots compare the changes in degradation rate constant upon SMG1 inhibition (x-axis) to the change in estimated isoform abundance upon SMG1 inhibition (y-axis). **Left:** points colored by density. **Right:** highlights three sets of isoforms: 1) those that are both strongly stabilized (L2FC($k_{deg}$) < -1) and strongly upregulated (L2FC(TPM) > 1); 2) those that are not strongly stabilized (L2FC($k_{deg}$) > -0.25) but are strongly upregulated; and 3) those that are strongly stabilized but not strongly upregulated (L2FC(TPM) < 0.25). **(C)** Extent to which isoform stability explains isoform abundance. **Left:** comparison of log-scale transcript isoform degradation rate constant in DMSO treated cells (x-axis) to its abundance (y-axis). Red line represents the linear best fit. **Middle:** comparison of the transcript isoform degradation rate constant (x-axis) to its isoform fraction (the fraction of transcripts from a gene contributed by a specific isoform) on a log-odds (logit) scale (y-axis). Red line is the linear best fit. **Right:** coefficient of determination for the linear regressions shown in the **Left** and **Middle** plots. Y-axis is thus interpretable as the fraction of isoform-to-isoform variance in either of the two y-axis quantities in those plots that is explained by the transcript isoform degradation rate constant.

### Figure S3

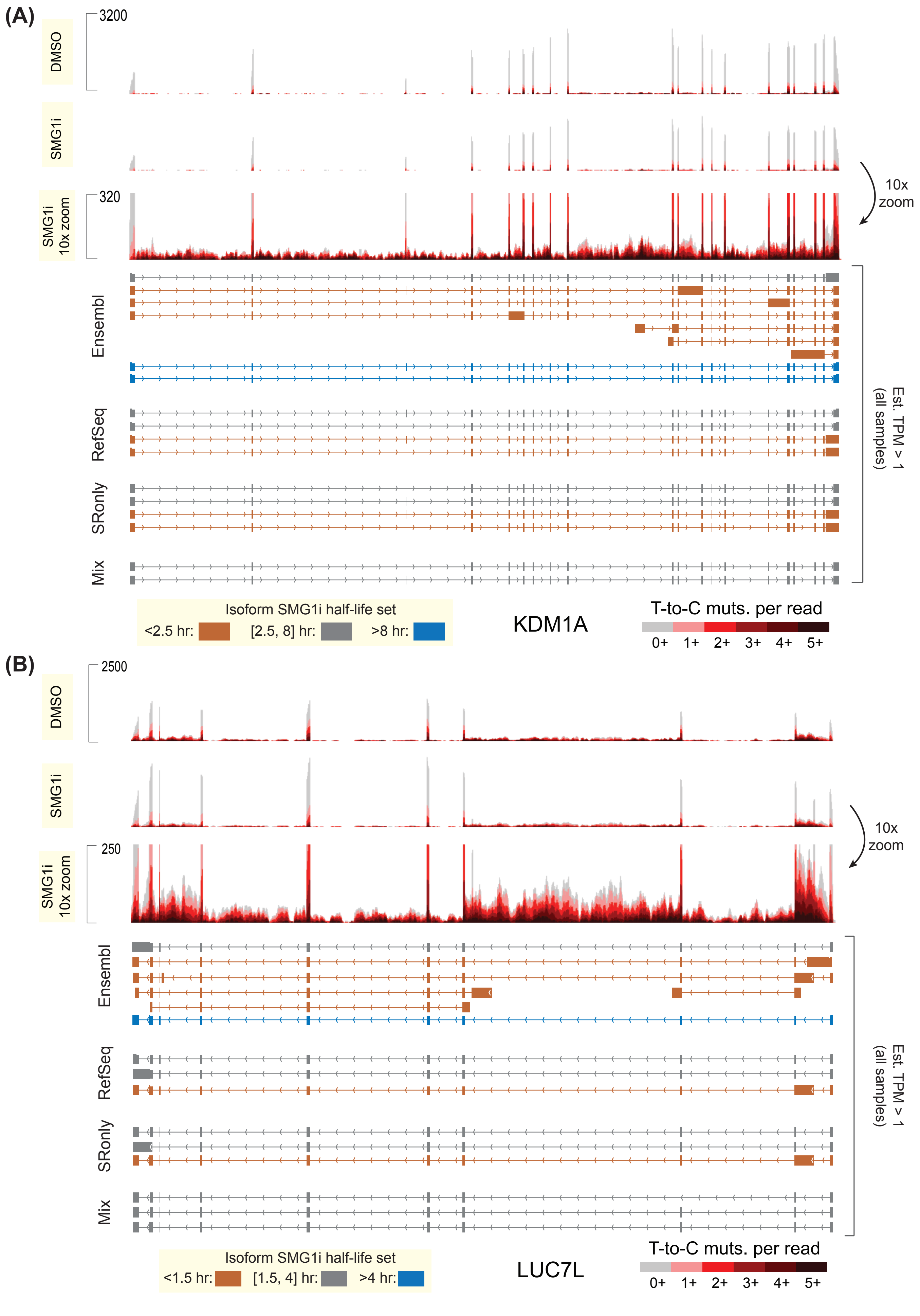

(A)
3200
DMSO
SMG1i
320
SMG1i 10x zoom
10x zoom
Ensembl
RefSeq
SRonly
Mix
Est. TPM > 1 (all samples)
Isoform SMG1i half-life set
<2.5 hr:
[2.5, 8] hr:
>8 hr:
KDM1A
T-to-C muts. per read
0+ 1+ 2+ 3+ 4+ 5+
(B)
2500
DMSO
SMG1i
250
SMG1i 10x zoom
10x zoom
Ensembl
RefSeq
SRonly
Mix
Est. TPM > 1 (all samples)
Isoform SMG1i half-life set
<1.5 hr:
[1.5, 4] hr:
>4 hr:
LUC7L
T-to-C muts. per read
0+ 1+ 2+ 3+ 4+ 5+

**Figure S3.** Associated with Figure 3
**(A)** TimeLapse-seq sequencing tracks for KDM1A and the 4 annotations discussed in this work. **Top track:** A representative replicate of DMSO treated data. **Middle track:** A representative replicate of SMG1i treated data. **Bottom track:** Middle track with an approximately 10x zoomed vertical scale to show intronic coverage and mutational content. **Annotations:** Isoform maps below tracks show isoforms with an RSEM estimated abundance of > 1 TPM in all samples. Isoforms are colored by their half-life, with isoforms binned into unstable (< 2.5 hour), moderately stable (between 2.5 and 8 hour), and highly stable (> 8 hour) transcripts. Note both the preponderance of highly unstable transcripts in the Ensembl annotation as well as the extent to which the presence of these isoforms inflates some estimates of stable isoform turnover.
**(B)** Same as **(A)** but for LUC7L. Transcript half-life set bounds are adjusted to better reflect the range of turnover kinetics at this locus.

Figure S4

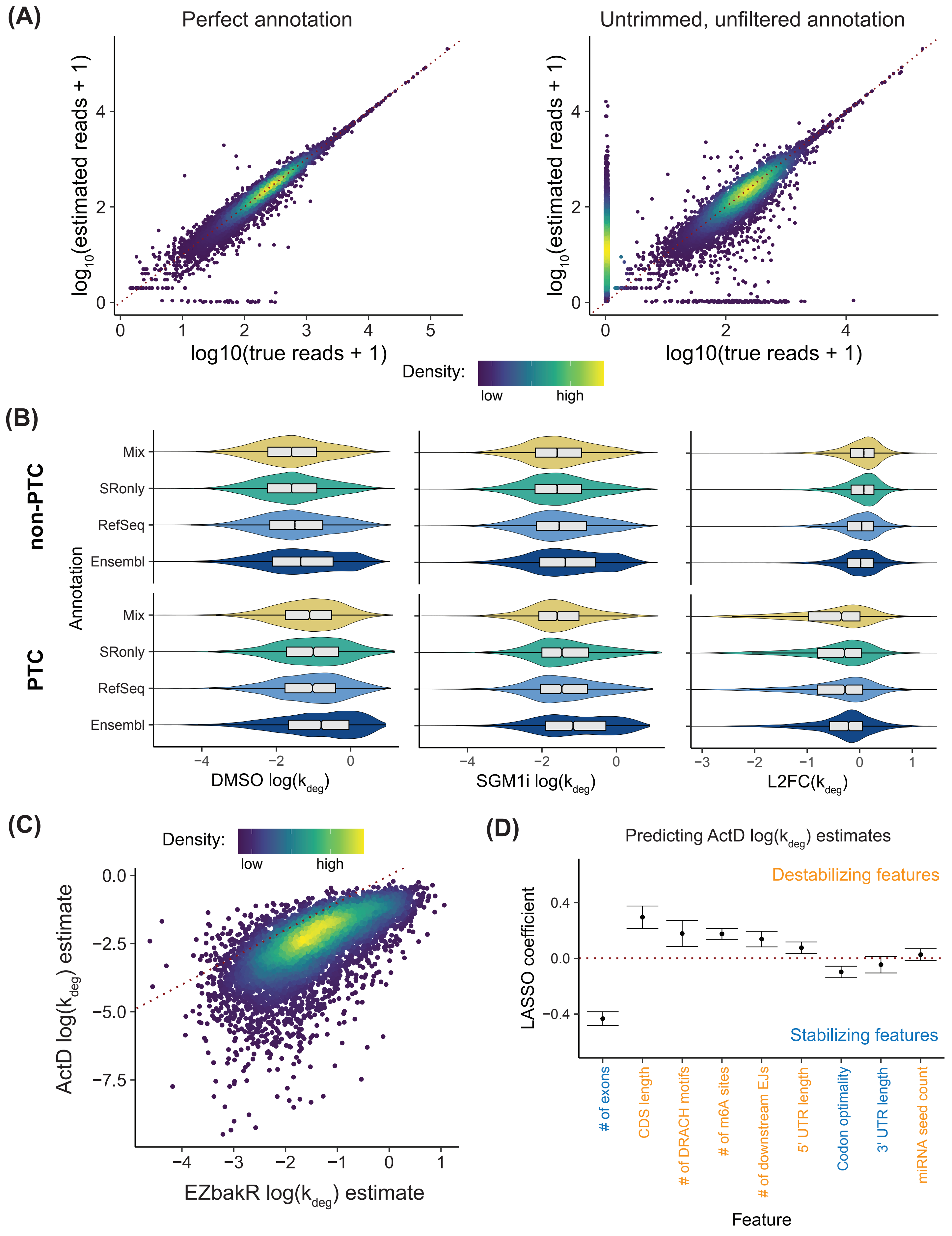

(A)
Perfect annotation
Untrimmed, unfiltered annotation
log10(estimated reads + 1)
log10(true reads + 1)
Density:
low
high
(B)
non-PTC
PTC
Annotation
Mix
SRonly
RefSeq
Ensembl
DMSO log(kdeg)
SGM1i log(kdeg)
L2FC(kdeg)
(C)
Density:
low
high
ActD log(kdeg) estimate
EZbakR log(kdeg) estimate
(D)
Predicting ActD log(kdeg) estimates
Destabilizing features
Stabilizing features
LASSO coefficient
### of exons
CDS length
### of DRACH motifs
### of m6A sites
### of downstream EJs
5' UTR length
Codon optimality
3' UTR length
miRNA seed count
Feature

**Figure S4.** Associated with Figure 3

**(A)** Assessment of transcript isoform quantification accuracy with simulated data and the use of different annotations. **Left:** Simulated data analyzed using the exact annotation from which the simulated data was generated. Same as what is used in **Figure 1** and **S1**. **Right:** Same simulated data but analyzed with an annotation including an amount of UTR overextension and number of unexpressed isoforms reflective of that seen in real data. Points are colored by density. Note the large number of reads spuriously assigned (points at x = 0) in simulations using the untrimmed, unfiltered annotation.

**(B)** Comparison of turnover kinetic trends seen in the 4 annotations assessed. **Left:** log-scale degradation rate constants in DMSO treated cells (x-axis) for all isoforms with estimated TPM > 1 in all samples. **Middle:** Same as **Left** but for SMG1i treated cells. **Right:** Changes in degradation rate constants upon SMG1i treatment for all isoforms with estimated TPM > 1 in all samples.

**(C)** Comparison of NR-seq/EZbakR derived isoform degradation rate constant estimates to those obtained from transcript isoform quantification of a previously published transcription inhibition RNA-seq time course, also in HEK293 cells [44]. Points are colored by density.

**(D)** LASSO regression coefficients for the same analysis shown in **Figure 3F**, but with log-scale degradation rate constants estimated from the transcription inhibition RNA-seq time course rather than our TimeLapse-seq data.

### Figure S5

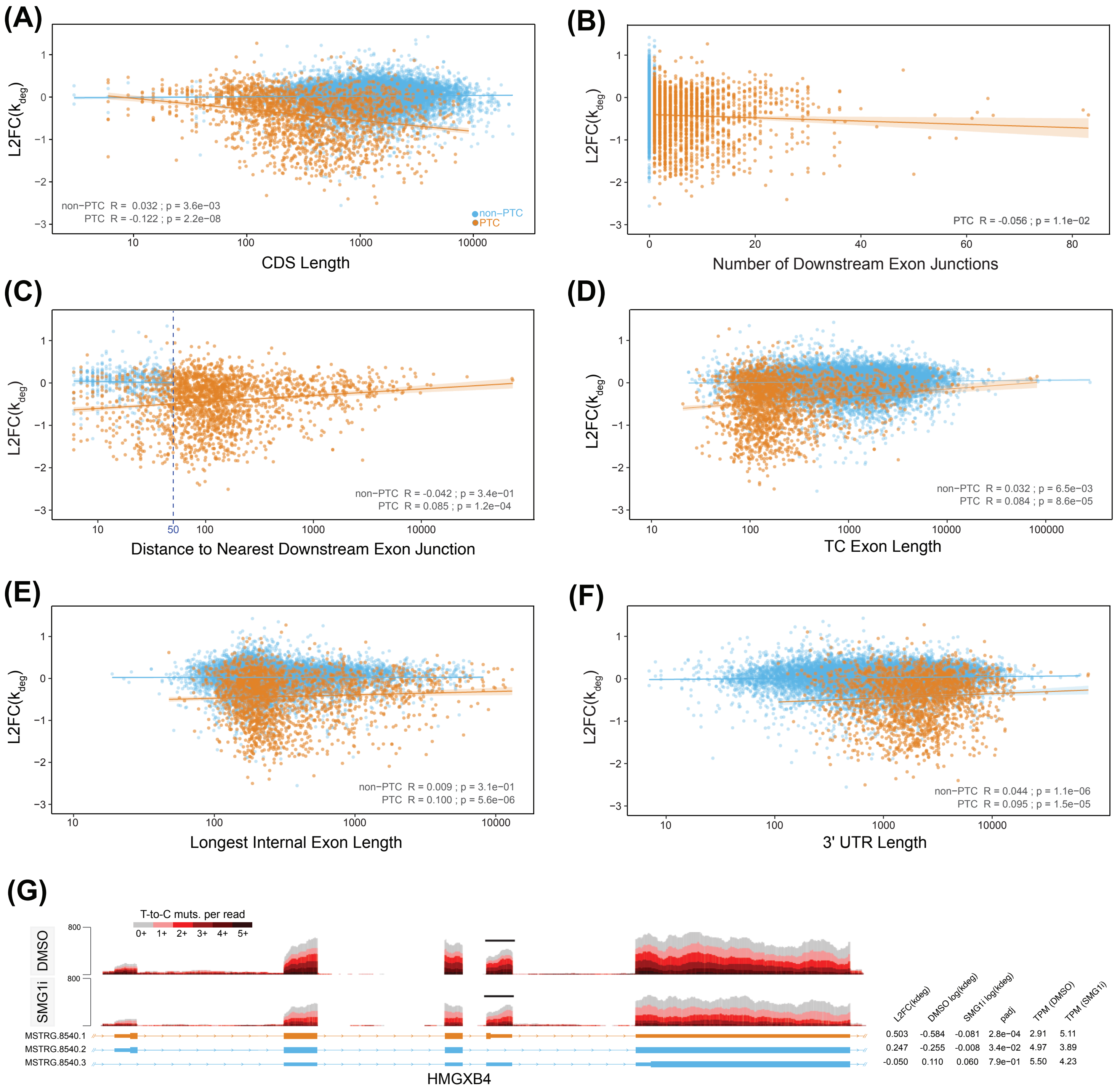

(A)
L2FC(k_deg)
non−PTC R = 0.032 ; p = 3.6e−03
PTC R = -0.122 ; p = 2.2e−08
non−PTC
PTC
CDS Length
(B)
PTC R = -0.056 ; p = 1.1e−02
Number of Downstream Exon Junctions
(C)
non−PTC R = -0.042 ; p = 3.4e−01
PTC R = 0.085 ; p = 1.2e−04
Distance to Nearest Downstream Exon Junction
(D)
non−PTC R = 0.032 ; p = 6.5e−03
PTC R = 0.084 ; p = 8.6e−05
TC Exon Length
(E)
non−PTC R = 0.009 ; p = 3.1e−01
PTC R = 0.100 ; p = 5.6e−06
Longest Internal Exon Length
(F)
non−PTC R = 0.044 ; p = 1.1e−06
PTC R = 0.095 ; p = 1.5e−05
3' UTR Length
(G)
T-to-C muts. per read
0+ 1+ 2+ 3+ 4+ 5+
DMSO
SMG1i
800
L2FC(kdeg) DMSO log(kdeg) SMG1i log(kdeg) padj TPM (DMSO) TPM (SMG1i)
MSTRG.8540.1 0.503 -0.584 -0.081 2.8e-04 2.91 5.11
MSTRG.8540.2 0.247 -0.255 -0.008 3.4e-02 4.97 3.89
MSTRG.8540.3 -0.050 0.110 0.060 7.9e-01 5.50 4.23
HMGXB4

**Figure S5.** Associated with Figure 5

(**A-F**) Scatter plots showing the $\log_2$-fold change in the isoform degradation rate constant (L2FC($k_{deg}$)) between SMG1i and DMSO treatments as a function of (**A**) CDS length, (**B**) number of downstream exon junctions, (**C**) distance to nearest downstream exon junction, (**D**) TC exon length, (**E**) longest internal exon length, and (**F**) 3'UTR length for non-PTC (blue) and PTC-containing (orange) isoforms. Linear regression fit lines with standard error shading are shown for PTC data, along with corresponding correlation coefficients (R) and significance values (p-value) for each regression. The vertical line at 50 nucleotides in (*C*) depicts the "50-55 nucleotide" rule.

(**G**) TimeLapse-seq tracks for the HMGXB4 gene under DMSO (upper tracks) and SMG1i (lower tracks) treatment conditions. The degree of $s^4U$ labeling is represented by increasing darkness of red, corresponding to the number of T-to-C mutations per read, with unlabeled reads shown in gray. Three HMGXB4 isoforms are schematized at the bottom, with the ORFs represented by thicker lines and the black bar indicating the position of the poison exon. Blue isoforms are non-PTC isoforms, the orange isoform is PTC-containing. Decay kinetics and RNA metrics for these isoforms are provided on the right.

### Figure S6

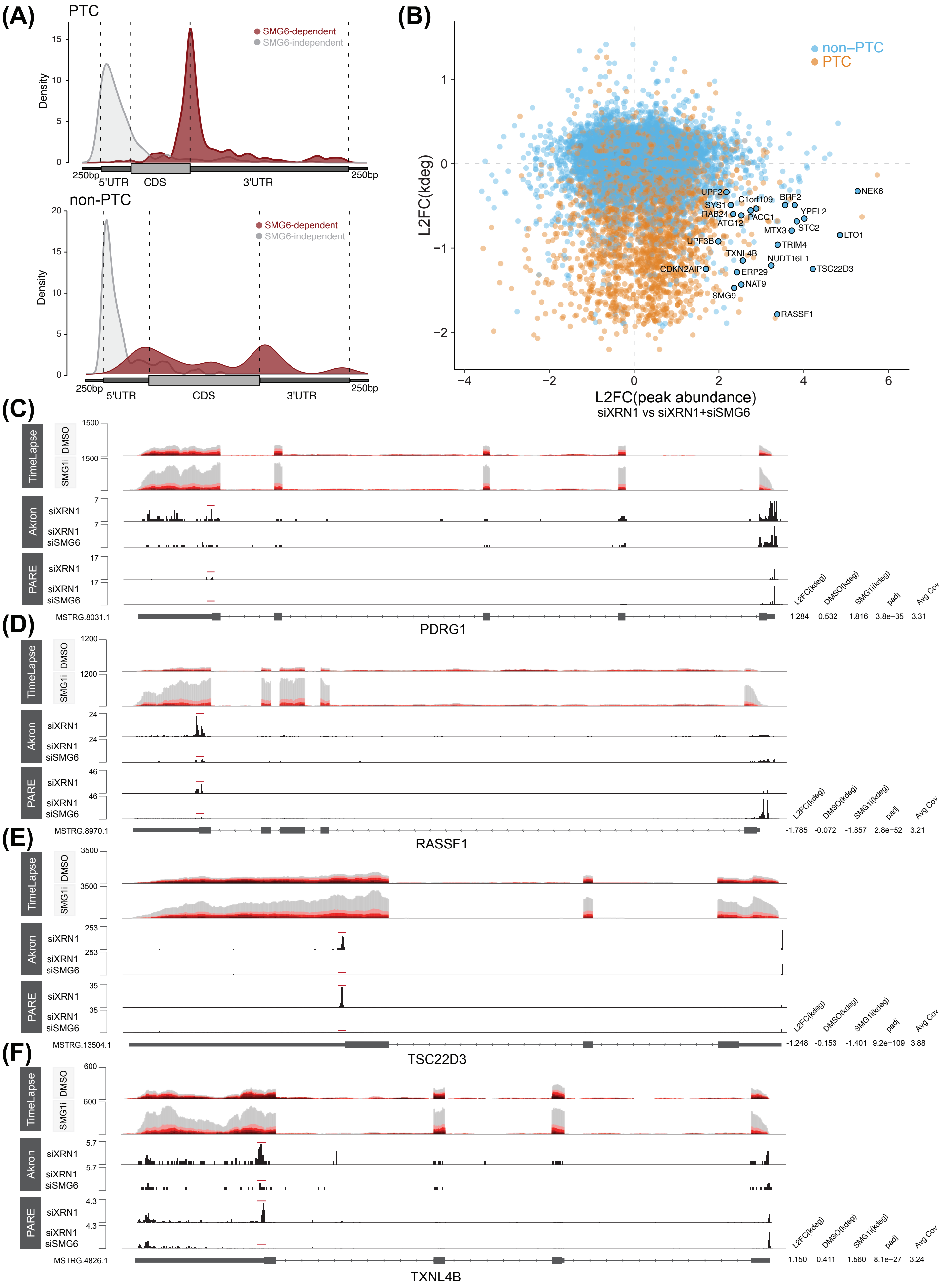

(A)
PTC
SMG6-dependent
SMG6-independent
Density
250bp
5'UTR
CDS
3'UTR
non-PTC
(B)
non–PTC
PTC
L2FC(kdeg)
L2FC(peak abundance)
siXRN1 vs siXRN1+siSMG6
NEK6
UPF2
SYS1
C1orf109
BRF2
YPEL2
RAB24
PACC1
ATG12
MTX3
STC2
LTO1
UPF3B
TRIM4
TXNL4B
NUDT16L1
TSC22D3
CDKN2AIP
ERP29
NAT9
SMG9
RASSF1
(C)
TimeLapse
DMSO
SMG1i
Akron
siXRN1
siXRN1 siSMG6
PARE
MSTRG.8031.1
PDRG1
L2FC(kdeg) DMSO(kdeg) SMG1i(kdeg) padj Avg Cov
-1.284 -0.532 -1.816 3.8e−35 3.31
(D)
MSTRG.8970.1
RASSF1
-1.785 -0.072 -1.857 2.8e−52 3.21
(E)
MSTRG.13504.1
TSC22D3
-1.248 -0.153 -1.401 9.2e−109 3.88
(F)
MSTRG.4826.1
TXNL4B
-1.150 -0.411 -1.560 8.1e−27 3.24

### Figure S6 continued

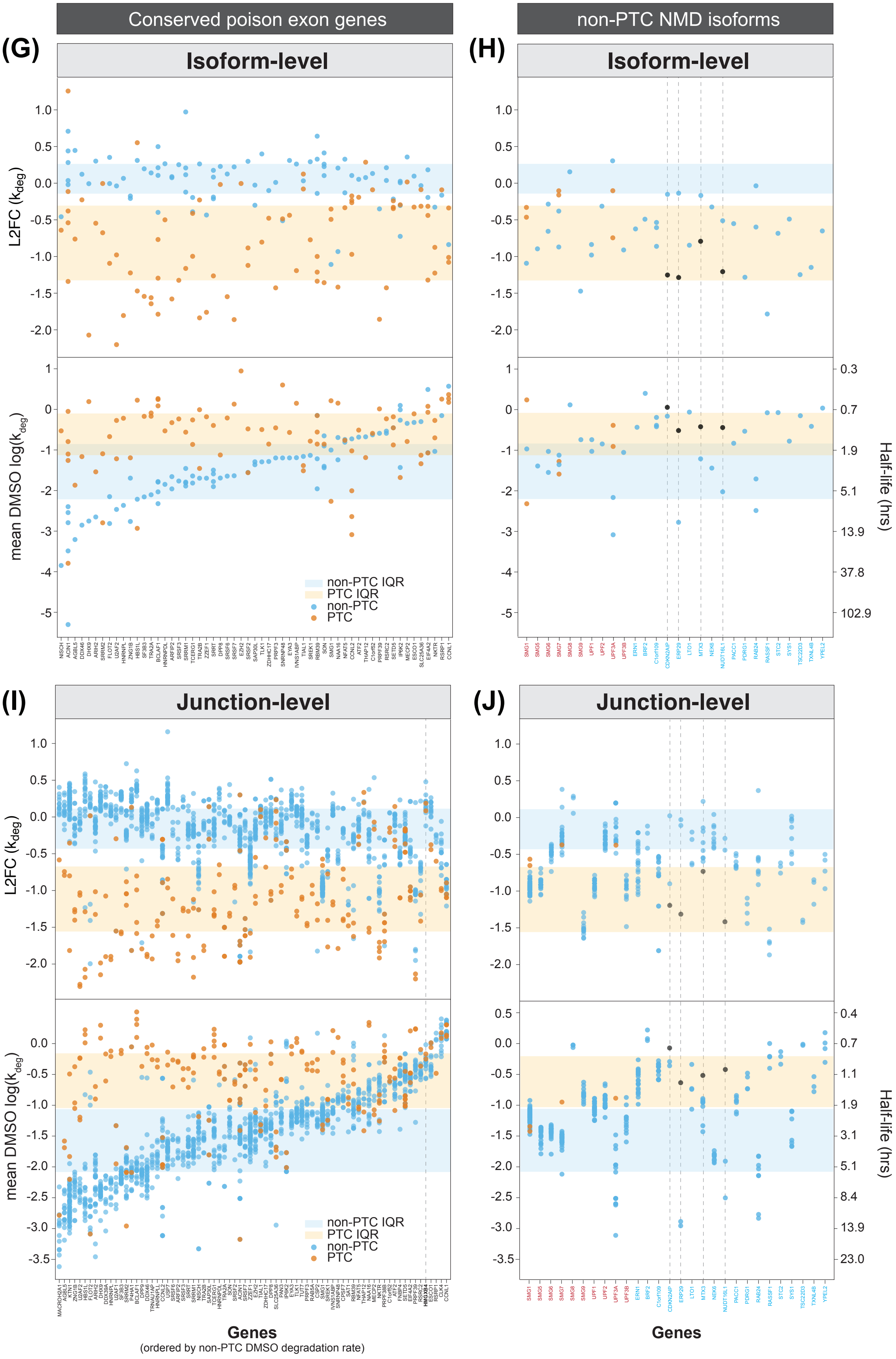

**Figure S6.** Associated with Figure 6

(**A**) Meta-gene analysis showing the distribution of peaks with significantly increasing (red) or decreasing (gray) L2FC(peak abundance) in siXRN1 vs. siXRN1+siSMG6 conditions. The top panel represents PTC-containing isoforms, while the bottom panel represents non-PTC isoforms. Below each distribution is a schematic of a meta-transcript. Gray peaks primarily correspond to increased decapping resulting from SMG6 knockdown, while red peaks indicate free 5' phosphates resulting from SMG6-dependent cleavage events.

**(B**) Scatter plot comparing isoform-level L2FC(peak abundance) between siXRN1 and siXRN1+siSMG6 conditions (x-axis) to L2FC($k_{deg}$) between SMG1i and DMSO treatments (y-axis) for non-PTC (blue) and PTC-containing (orange) isoforms. Genes highlighted in the figure are discussed further in the text and throughout Figure S6.

(**C-F**) **Top:** TimeLapse-seq tracks for PDRG1 (**C**), RASSF1 (**D**), TSC22D3 (**E**) and TXNL4B (**F**) under DMSO (upper tracks) and SMG1i (lower tracks) treatment conditions. The degree of $s^4U$ labeling is indicated by increasing darkness of red, corresponding to the number of T-to-C mutations per read, with unlabeled reads shown in gray. **Bottom:** Akron-seq (this study) and PARE-seq [56] tracks from XRN1 and XRN1+SMG6 depletion experiments, highlighting SMG6-dependent cleavage sites marked by red lines above the sequence tracks. The isoform from each gene is schematized below, with the ORFs represented by thicker lines. Corresponding decay rate measurements for these isoforms are displayed to the right.

(**G**) Distribution of isoform-level stability measurements for conserved poison exon-containing genes. **Top:** Dot plots displaying the distribution of isoform-level L2FC($k_{deg}$) changes between SMG1i and DMSO treatments for a set of conserved poison exon-containing genes. **Bottom:** Dot plots showing the distribution of isoform-level DMSO log($k_{deg}$) decay rate measurements for the same set of genes. Genes are ordered by the DMSO degradation rate of their non-PTC isoforms, from lowest to highest. Blue dots represent non-PTC isoforms, and orange dots represent PTC-containing isoforms. The orange shaded region highlights the interquartile range (IQR) of L2FC($k_{deg}$) changes in SMG1i vs. DMSO (*top*) or the mean DMSO $k_{deg}$ values (bottom) for PTC-containing isoforms. Similarly, the blue shaded region indicates the IQR of L2FC($k_{deg}$) changes in SMG1i vs. DMSO for non-PTC isoforms.

(**H**) As in (G), but for non-PTC isoforms. The shaded regions highlight the IQRs of the L2FC($k_{deg}$) in SMG1i vs. DMSO (*top*) or the mean DMSO $k_{deg}$ values (bottom) for PTC-containing isoforms (orange) and nonPTC isoforms (blue) of poison exon-containing genes (same ranges as in G). Genes are color-coded, with NMD factors in red and high-confidence non-PTC isoforms in blue.

(**I-J**) Junction-based stability measurements for validated PTC **(I)** and non-PTC **(J)** NMD-targeted genes. **Top:** Dot plots showing the distribution of junction-level L2FC($k_{deg}$) in degradation rate measurements between SMG1i and DMSO treatments. **Bottom:** Dot plots showing the distribution of junction-level DMSO log($k_{deg}$) degradation rate measurements for the same set of genes, ordered and colored as in (G) and (H). Blue dots represent non-PTC junctions, while orange dots represent junctions associated with PTC-exons. The orange shaded region highlights the IQR of L2FC($k_{deg}$) changes in SMG1i vs. DMSO (*top*) or the mean DMSO log($k_{deg}$) values (*bottom*) for PTC-containing isoforms of poison exon-containing genes. Similarly, the blue shaded region indicates the IQR of L2FC($k_{deg}$) changes in SMG1i vs. DMSO for non-PTC isoforms of poison exon-containing genes

### Figure S7

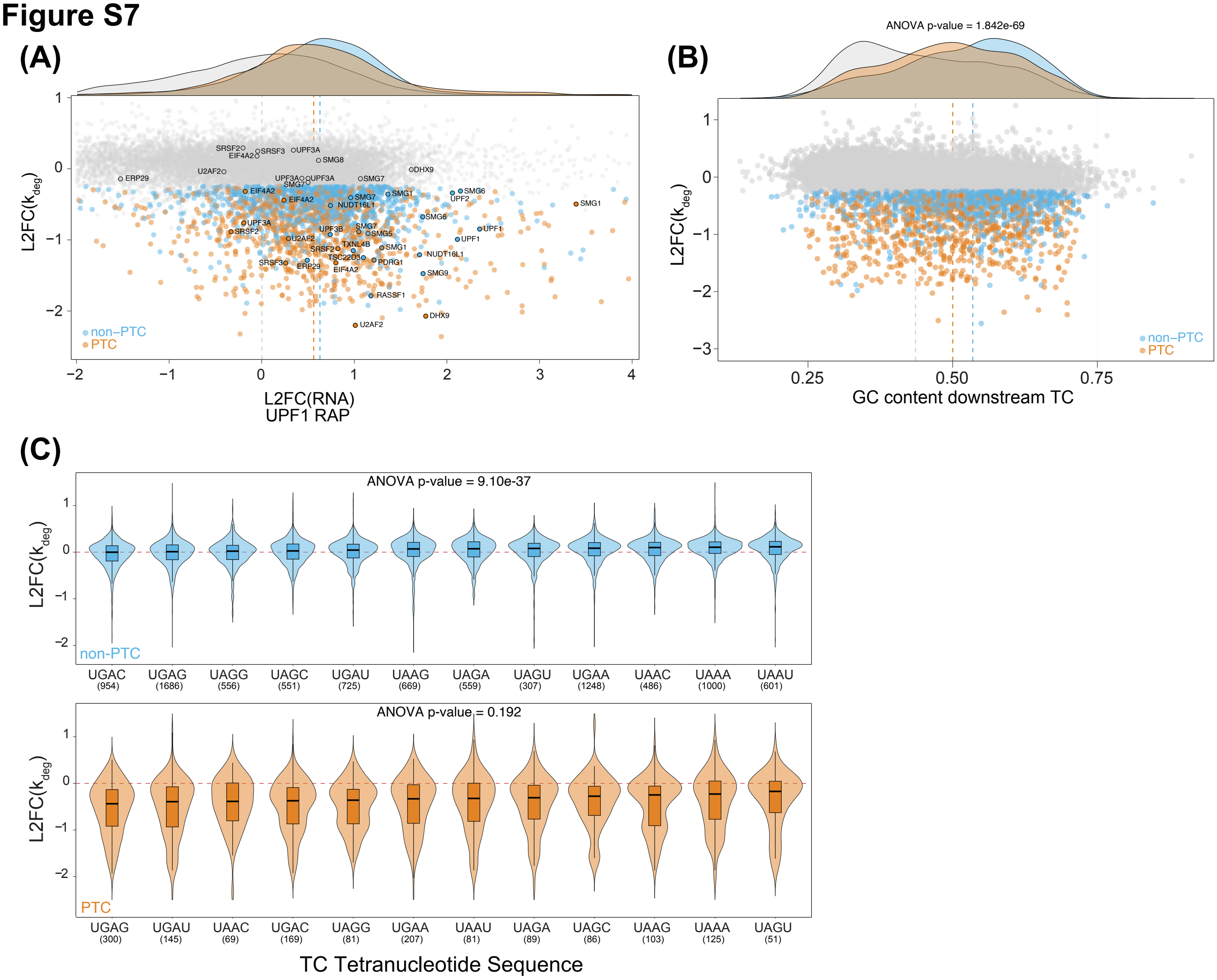

(A)
L2FC(k_deg)
non–PTC
PTC
L2FC(RNA)
UPF1 RAP
(B)
ANOVA p-value = 1.842e-69
L2FC(k_deg)
non–PTC
PTC
GC content downstream TC
(C)
ANOVA p-value = 9.10e-37
L2FC(k_deg)
non-PTC
UGAC (954)
UGAG (1686)
UAGG (556)
UAGC (551)
UGAU (725)
UAAG (669)
UAGA (559)
UAGU (307)
UGAA (1248)
UAAC (486)
UAAA (1000)
UAAU (601)
ANOVA p-value = 0.192
L2FC(k_deg)
PTC
UGAG (300)
UGAU (145)
UAAC (69)
UGAC (169)
UAGG (81)
UGAA (207)
UAAU (81)
UAGA (89)
UAGC (86)
UAAG (103)
UAAA (125)
UAGU (51)
TC Tetranucleotide Sequence

### Figure S7 continued

## (D)

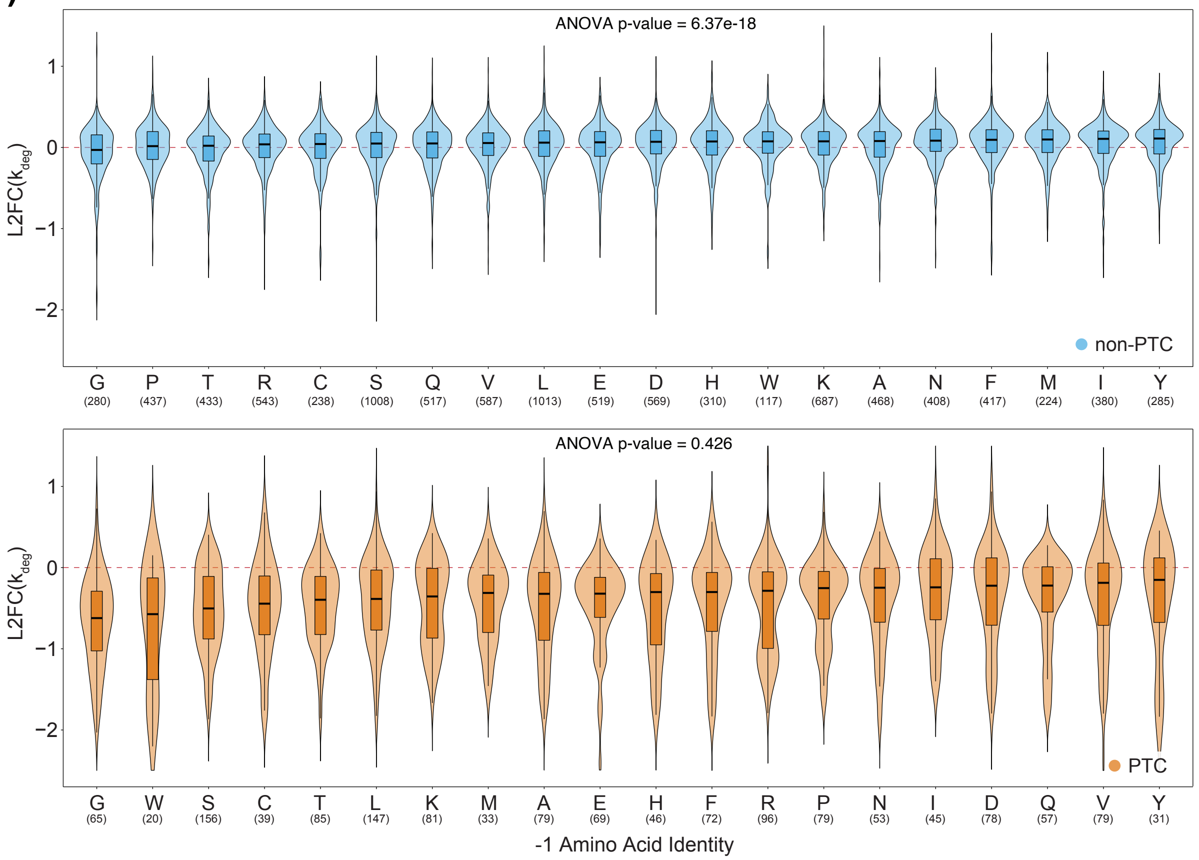


## (E)

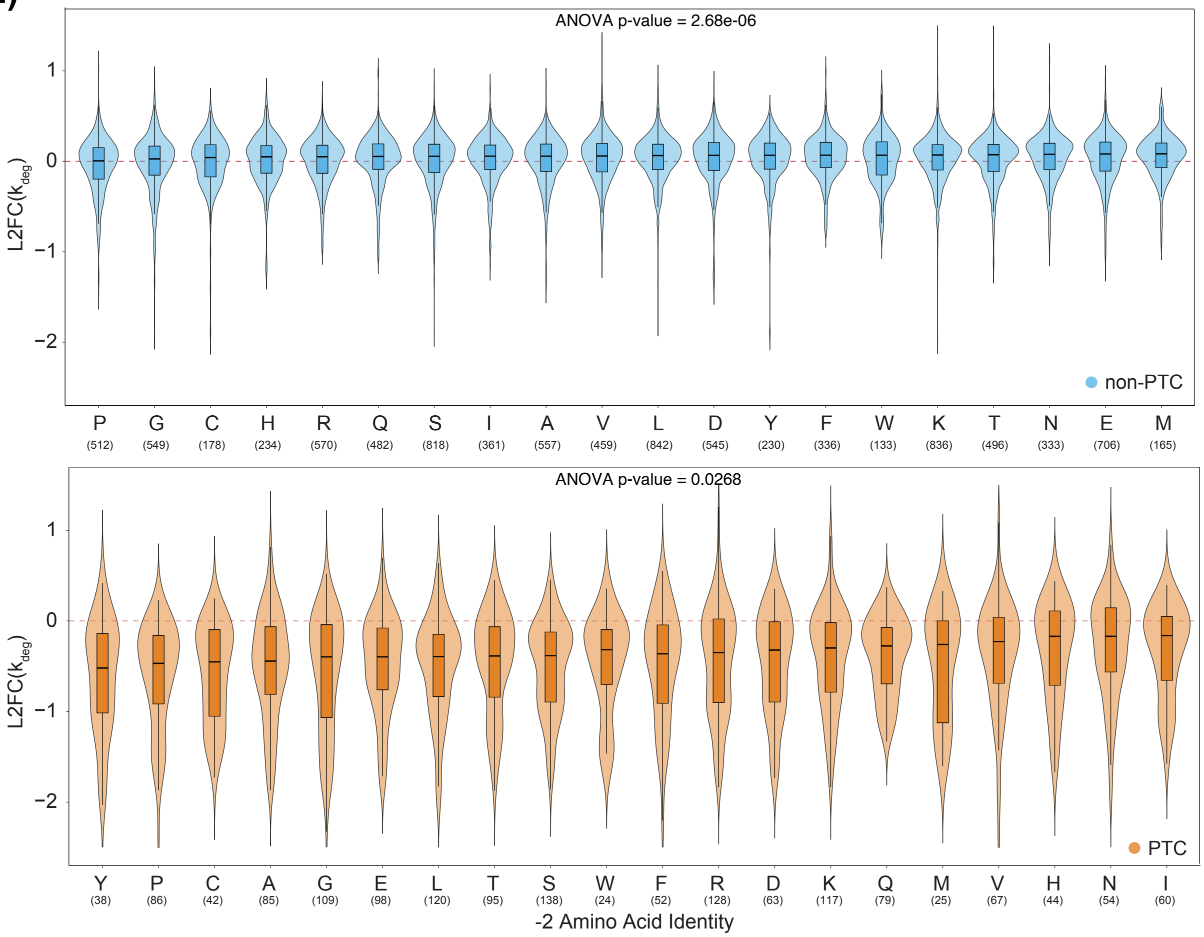

**Figure S7.** Associated with Figure 7

(**A**) Scatter plot showing L2FC($k_{deg}$) differences between SMG1i and DMSO treatments versus L2FC(RNA) enrichment from CLIP-UPF1 RNA affinity purification (RAP). SMG1i-responsive non-PTC and PTC isoforms are represented in blue and orange, respectively, and non-responsive isoforms are in gray. Vertical lines indicate the median of each group's distribution. Data are the same as Figure 7C and isoforms are labeled as in Figure 6. **Top:** Density plot of each group with vertical lines indicating the median of each group's distribution.

(**B**) Scatter plot of L2FC($k_{deg}$) versus GC-content 200 nt downstream of TC, colored as in (A). **Top:** Density plot for each group, with vertical lines indicating the median of each distribution. ANOVA p-value reflects statistical significance as described in **Figure 7D**. (**C-E**) Violin plots showing L2FC($k_{deg}$) differences between SMG1i and DMSO treatments for non-PTC (blue, *top*) and PTC (orange, *bottom*) isoforms, grouped by (**C**) TC tetranucleotide sequence, (**D**) C-terminal -1 amino acid, and (**E**) C-terminal -2 amino acid. The horizontal dashed red line represents no change in L2FC($k_{deg}$) between conditions. Sample sizes for each category are shown below the plots. ANOVA p-value reflects statistical significance as described in **Figure 7D**.
